# Target-matched increased fidelity SpCas9 variants can edit any target without genome-wide off-target effects

**DOI:** 10.1101/2022.08.16.504097

**Authors:** Péter István Kulcsár, András Tálas, Zoltán Ligeti, Eszter Tóth, Zsófia Rakvács, Vanessza Laura Végi, Sarah Laura Krausz, Krisztina Huszár, Ervin Welker

**Affiliations:** Institute of Enzymology, Research Centre for Natural Sciences, Budapest, Hungary; Institute of Biochemistry, Biological Research Centre, Szeged, Hungary; Doctoral School of Multidisciplinary Medical Science, University of Szeged, Hungary; Biospiral-2006 Ltd, Szeged, Hungary; School of Ph.D. Studies, Semmelweis University, Budapest, Hungary; Gene Design Ltd, Szeged, Hungary

## Abstract

*Streptococcus pyogenes* Cas9 (SpCas9) has been employed as a genome engineering tool with a promising potential within therapeutics. However, its off-target effects pose a major limitation for applications requiring high specificity. Approaches developed to date to mitigate this effect, including any of the increased fidelity (i.e., high-fidelity) SpCas9 variants, only provide efficient editing on a relatively small fraction of targets without detectable off-targets. Upon addressing this problem, we have revealed a rather unexpected cleavability ranking of target sequences, and a cleavage rule that governs the on-target and off-target cleavage of increased fidelity SpCas9 variants but not that of SpCas9-NG or xCas9. As a consequence of this rule, each target needs a matching-fidelity, optimal variant for efficient cleavage without detectable off-target effects. By exploiting this finding, we have developed an extended set of increased fidelity variants spanning across a wide range, with differences in fidelity small enough to comprise an optimal variant for any target, irrespective of its cleavability ranking. We have demonstrated efficient editing with maximum specificity even on those targets that have been challenged but failed in previous studies.

## Introduction

Although many challenges remain to be addressed until advances of the CRISPR technology can be translated into routine clinical practice, recent reports on both *in vivo* and *ex vivo* CRISPR-based gene therapy reaching the stage of clinical trials mark the enormous potential of CRISPR nucleases^1–10^. The *Streptococcus pyogenes* Cas9 (SpCas9) is the most frequently used nuclease for genome engineering with the highest potential for therapeutic applications amongst all RNA-guided nucleases of the type II CRISPR system. Tremendous research effort has been devoted to increase the potential of SpCas9 by minimizing its off-target activity, which poses safety concerns for its use in areas where high specificity is a requirement, e.g., in clinical applications^4, 11–13^. Several methods have been developed to increase its specificity including the application of double nickases^14, 15^ and dimer FokI fusion variants of SpCas9^16–18^, single-guide RNAs (sgRNAs) with truncated or extended spacers^18–21^, as well as mutant SpCas9 variants^4^. However, none of these have managed to fully eliminate off-target cleavage and/or preserve efficient on-target editing universally for most targets. One of the most promising approaches among these methods to decrease off-target activity has been the generation of increased fidelity nuclease variants. A non-exhaustive list of these variants include the rationally designed eSpCas9, SpCas9- HF1, HypaSpCas9 and Blackjack^21–26^, as well as variants developed by a selection scheme; evoSpCas9, Sniper, and HiFi^27–31^. A number of variants have also been developed by combining mutations from existing increased fidelity nucleases (IFNs), these include e-plus, HF1-plus, HypaR, and HeFSpCas9^21, 24, 25, 32^. We prefer to collectively refer to the variants as ‘increased fidelity’ nucleases instead of ‘high fidelity’ nucleases, because the term ‘high-fidelity’ have been reserved specifically for the SpCas9-HF variants^23^, and also because, as it will be clear from this paper, they possess widely varying fidelities. While increased fidelity variants greatly improve the potential for highly specific genome modifications, their limitations have also become increasingly apparent. Each of them initiate the editing of many targets with considerable off-target effects^21–29, 31, 33^ and they exhibit increased target-selectivity, i.e., the variants do not initiate editing or they only do to a decreased extent at numerous target sites that are otherwise cleavable by the wild type (WT) SpCas9^21, 23–25, 33–35^. Our former *in cellulo* study also revealed that HeFSpCas9, one of the highest fidelity variants, cleaves a few targets only, albeit with high fidelity, however, these exact targets are the ones, that get cleaved by the enhanced- and SpCas9-HF1 variants with the most concomitant off-target effects^21, 33^. This finding prompted us to investigate whether this pattern is also a characteristic of other increased fidelity variants and target sequences. In this study we demonstrate that (i) IFNs fall into an order according to their fidelity/target-selectivity. Even more interestingly, we also show that different target sequences have substantially different effects on the activity of IFNs, and ultimately these sequence contributions control whether an IFN cleaves a target or not, and they also primarily determine the extent of their actual off-target propensity. For optimal, both highly efficient and specific editing, one should find an IFN with a fidelity/target selectivity ranking that is well matched to the sequence contribution of the target, i.e., the variant should have an activity that is sufficient to efficiently cleave the target sequence but insufficient to cleave any of its off-target sequences. (ii) The fidelity requirement of the potential target sequences is frequently not accounted for by the available variants. Therefore, to provide a near-optimal variant for any potential target, we generated new variants to build a set of IFNs with increasing fidelity with small enough differences between the variants to cover a wide range in high resolution. (iii) Using this knowledge and an extended set of variants, we show that practically every target can be edited without detectable genome-wide off-target effects (defined here as detectable by GUIDE-seq), by applying target-matched IFNs. We challenged this claim by testing all known problematic target sites from the literature that have been unsuccessfully tried by the previously developed, commonly used SpCas9 IFNs ^20, 23, 24, 27, 29^, as off- target editing was still detected by GUIDE-seq.

## Results

### Cleavage rule controls the on-target activity of increased fidelity nucleases

First, using an EGFP disruption assay in N2a cells, we compared the on-target activity of WT and seven IFNs; Blackjack-, e-plus, HF1-plus, Hypa-, HypaR-, evo- and HeFSpCas9 on 50 targets (target sequences can be found in Supplementary Table 1) using flow cytometry (gating examples in Supplementary Fig. 1, results in Supplementary Fig. 2a-i)^21, 24, 27, 32, 33^. The results are also presented on a heatmap depicting disruption activities for each target, normalized to the wild type value in order to neutralize the effect of the cellular context and factors, such as sgRNA expression levels and sequence specificity of the NHEJ DNA repair system (Supplementary Fig. 2j). The variants exhibited varying normalized average on-target activity on these targets, Blackjack SpCas9 showing the highest, approaching that of the wild type, and HeFSpCas9 showing the lowest. We found that the cleavage pattern is far from being random. By reordering variants based on the number of targets they cleave (Supplementary Fig. 2k), we noticed that, generally, when a target was cleaved by a nuclease, it was also cleaved by all other lower-ranking nucleases (i.e.: by all those variants that, in aggregate, cleave a larger number of targets in the set). Moreover, when we reordered the targets based on the number of variants that could cleave them (Supplementary Fig. 2l), we realized that, generally, when a nuclease cleaves a particular target, it also cleaves all other targets that are in higher position in the cleavability ranking. We named this phenomenon the *cleavage rule* of the targets and variants. It creates a particularly striking pattern, separating the cleaved and not-cleaved values into two distinct classes in the two-dimensional cleavage map (Supplementary Fig. 3a). Binary classification confirms that the actual data of the two-dimensional cleavage map in Supplementary Figure 3a tightly follow this cleavage rule (G-mean score of 0.987, for details see Methods section and Supplementary Table 4) containing hardly any outliers. ROC curve (receiver operating characteristic curve) shows the fitting of the cleavage data of each variant to the two-dimensional cleavage map arranged according to the cleavage rule in Supplementary Figure 3a, confirming that this rule applies for each variant (Supplementary Figure 3b).

### On-target and off-target activities of IFNs on a given target are interconnected

To find out how this cleavability characteristic of the sequences is related to the off-target propensities of the variants, we conducted a mismatch screen. In this, we tested three PAM-distal positions each containing all three possible single mismatches for each of the eighteen selected target sequences altogether with 162 mismatching sgRNAs (Supplementary Fig. 3c). In the case of all the IFNs, the specificity of the editing on a given target clearly depends on the position of the target within this ranking. The fidelity-increasing mutations in a given variant may reduce the activity of SpCas9 appropriately for a relatively small fraction of the target sequences, so that it cleaves the on-target sequences efficiently and exclusively, without cleaving the off-targets. This can be illustrated by the example of HypaSpCas9. Efficient cleavage with maximum specificity can be seen at targets 8, 15 and 34. However, target sequences from lower cleavability ranks, such as targets 7, 11 and 35, will not be cleaved at all and targets from higher cleavability ranks, such as targets 2, 3 and 5, are cleaved but with off-targets (Supplementary Fig. 3c).

Taken together, these results suggest that there are 3 main factors that combine to determine whether an IFN will cleave a target or its off-targets. Figure 1 shows how the sum of the effects of these three factors, namely sequence contribution, specific mismatch(es) and fidelity enhancing mutations affect SpCas9. There are substantially larger differences in the effect of sequence contributions of different targets than the effect of some off-target mismatches, thus no single IFN is capable of the off-target-free cleavage of all sequences. The cleavage rule also imposes that in order to efficiently edit a target with the highest possible specificity, we need to select the IFN with the highest fidelity that still yields sufficient cleavage required for the given application. Supplementary Figure 3d shows that by exploiting the cleavage rule we can substantially increase the specificity of efficient IFN editing, however, several targets are still edited with considerable off-target effects (20-70% of the on-target disruption values).

**Figure 1.**
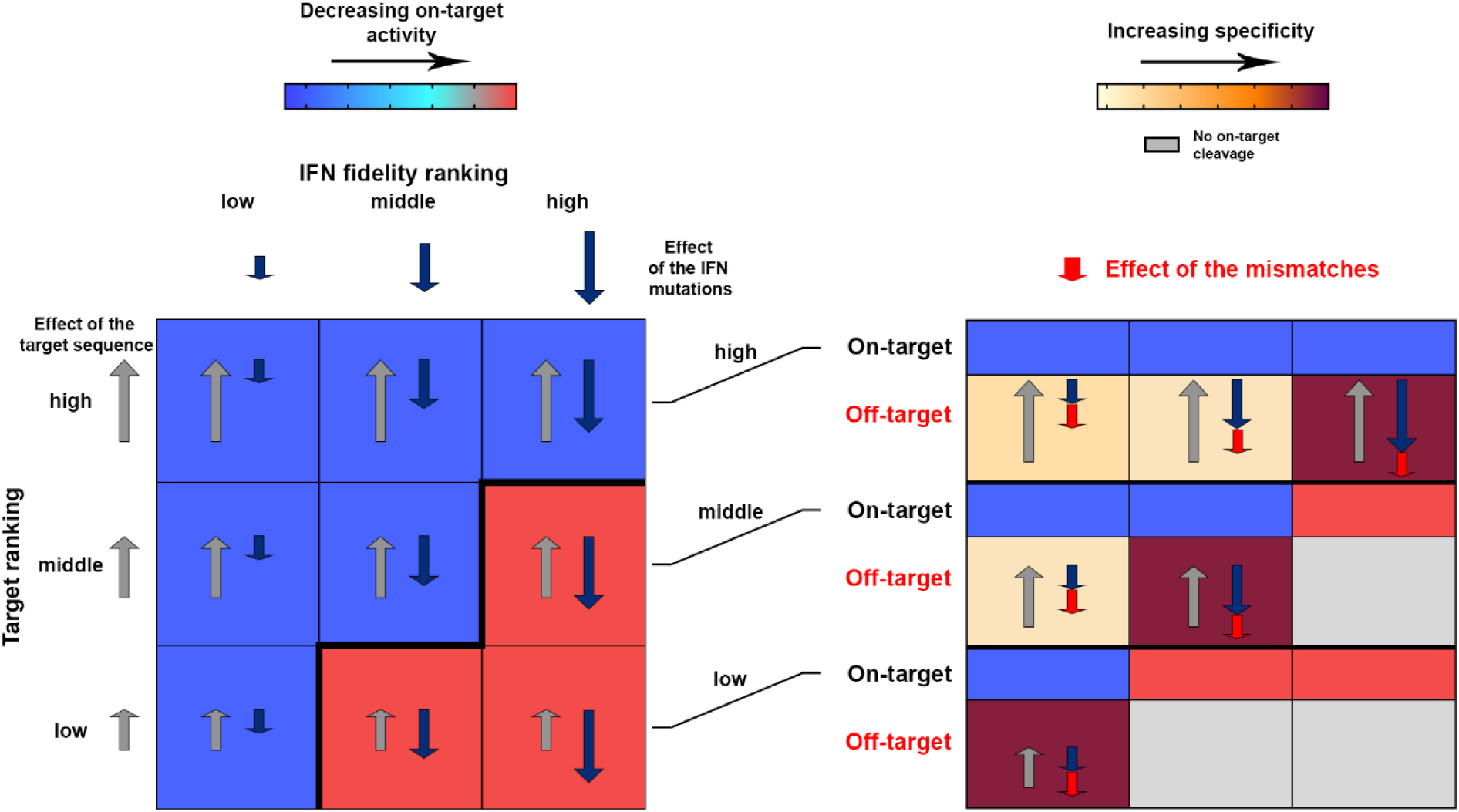
Simplified explanatory figure of the "cleavage rule". A simplified explanatory figure of the "cleavage rule" is presented to interpret the results shown in Figure 2 and Supplementary Figures 1 and 2. Three main factors, namely fidelity-increasing mutations, target sequence and mismatches combine to determine whether an IFN cleaves on-target and off-target sequences efficiently or not. The coloring of the heatmap in this figure corresponds to the coloring of the heatmaps shown in other figures in the manuscript. The left panel shows the activity of on-target IFNs of three different fidelity ranks on three targets of three different cleavability ranks: when the effect of sequence contribution is larger than the effect of fidelity-increasing mutations, the SpCas9 variant cleaves the target (blue background); when the effect is smaller, it does not cleave (red background). The right panel shows the effect of a mismatch on the cleavage of IFNs from the first panel on the same targets. When the effect of sequence contribution is larger than the combined effect of fidelity-increasing mutations and mismatches, the SpCas9 variant cleaves the off-target sequence (yellow background); when it is smaller, it does not cleave (brown background). The effect of the mutations in the optimal target-matched IFN is only slightly smaller than the contribution of the target sequence, and it effectively cleaves the on-target sequence. However, when the effect of any mismatches is added, the combined effect exceeds the contribution of the target sequence, and it does not cleave off-targets. The effect of the same mismatch may be different in different sequential contexts (https://www.biorxiv.org/content/10.1101/2021.11.18.469088v2). For simplicity, here we assign the same effect in all cases.

### Building a large set of IFNs with appropriate fidelities

We hypothesize that maximal fidelity can be achieved universally for every target sequence by having an extended set of IFNs with increasing fidelity. These IFNs should cover a wide range of fidelity rate in a sufficient resolution to provide an appropriate variant for targets from any cleavability ranks. To test this idea, we made use of our prior discovery that Blackjack mutations in SpCas9 variants not only make the 5’G extension of sgRNAs more tolerable, but they also increase their fidelity to some extent^21^. By generating new variants we established a set of 19 IFNs in total, including Sniper, HiFi, e-, -HF1, Hypa-, HypaR-, evo-, HeF-, their Blackjack counterparts (indicated with a ‘B’ prefix), e-plus, HF1-plus and Blackjack SpCas9^21–23, 28, 29, 32^. All variants expressed at similar levels as demonstrated by western blot (Supplementary Fig. 4). We found that all newly added variants fit in the pattern seen in Supplementary Figure 2 when tested on the on-target and mismatch screens (Fig. 2). Containing hardly any outliers (G-mean of 0.984) they all strictly follow the cleavage rule (Fig. 2a). When the highest fidelity variant with sufficient activity is used for each target (Fig. 2e), the specificity of IFN-editing is substantially increased. Furthermore, overall a higher specificity could be reached using this set than with the set of only 7 IFNs seen earlier in Supplementary Figure 2d (Fig. 2e). The 20-70% normalised off-target edits seen in Supplementary Figure 2d are effectively diminished by using this set of 19 variants suggesting that they approximate an appropriate resolution (Fig. 2e). The SpCas9 variants with the highest fidelity rank from our set of 19 IFNs that still show sufficient activity on a given target are hereafter referred to as target-matched variants for that given target. These results suggest that maximal fidelity can be reached universally by using an appropriate set of SpCas9 variants with small enough differences in fidelity that can provide an optimal target-matched IFN to every target from any position of the cleavability ranking.

**Figure 2.**
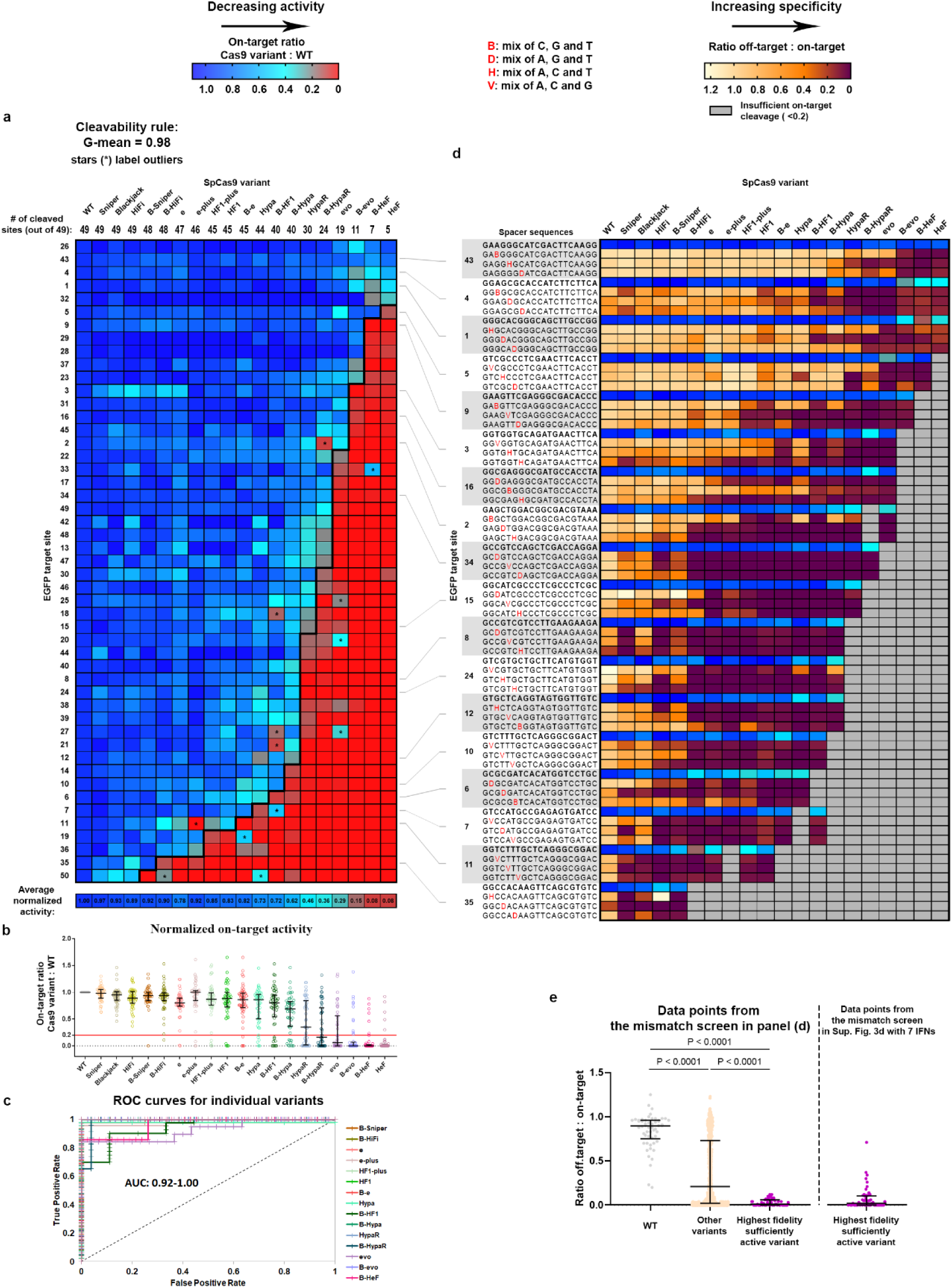
Extending the set of IFNs enables increased specificity. Heatmaps show the normalized EGFP disruption activity of SpCas9 nucleases with (**a**) perfectly matching and (**d**) partially mismatching 20G-sgRNAs in N2a cells. **a**, The bold line indicates the dividing line defined by the cleavage rule between the classes of cleaved and not-cleaved values. The G-mean value indicates how well the data points above and below the bold line correspond to cleaved and not-cleaved (<0.20 activity normalized to WT) experimental values. Targets and IFNs are shown in the same order as in Supplementary Figure 3. **b,** Normalized on-target disruption activities of various SpCas9 variants presented on a scatter dot plot. The sample points correspond to data presented in panel (**a**). The median and interquartile range are shown; data points are plotted as open circles representing the mean of biologically independent triplicates. Continuous red line indicates 0.20 normalized disruption activity, under which we consider the IFNs not to be active on a given target. Statistical significance was assessed by using RM one-way ANOVA and is shown in Supplementary Table 9. **c,** The ROC curves demonstrate that the order of the target sequences, determined by the cleavage rule, competently separates the classes of cleaved and not-cleaved normalized disruption values of each of the 19 variants from panel (**a**). **d**, Targets from higher ranks (cleavable by many IFNs) require higher fidelity nucleases, while targets from lower ranks (cleavable by few IFNs) require lower fidelity nucleases for editing with both high efficiency and high specificity. **e,** Matching IFNs and targets further increases the specificity of editing. The median and interquartile range of data points that are selected from panel (**d**) is presented as indicated; n=54, 654, 54, respectively. Dots are shown for each variant with each mismatching spacer position, provided that the on-target activity exceeded 70%; data are omitted otherwise. **a-e,** Target sequences, raw and processed disruption data and statistical details are reported in Supplementary Tables 1-4 and 9.

### The cleavage rule appears to be universal

To validate our findings, (i) we assessed the mismatch tolerance by genome-wide off-target detection instead of a mismatch screen, (ii) tested another cell line and used NGS instead of a disruption assay and (iii) analysed data from a large target library, as described below. (i) To validate that the characteristics of IFNs revealed by mismatch screening reliably reflect their genome-wide off-target effects, we performed GUIDE-seq analyses using various IFNs on 4 EGFP targets from different cleavability ranks of Figure 2 in HEK293.EGFP cell line. Figure 3 (and Supplementary Fig. 5) shows that the genome-wide off-target effects of an IFN correspond to its EGFP assay mismatch tolerance, both being primarily determined by the position of the target and the IFN within the two-dimensional ranking map. The number of off-targets in Guide-seq and the specificity of editing in the disruption assay change in parallel: decrease and increase, respectively. (ii) The fidelity rank of the IFNs and the cleavage rule remained in effect when cleavage was examined in different cell line (HEK293) and on 52 endogenous target sites (G-mean= 1.00, Fig. 4a-b). (iii) We further verified these results on the largest possible target data set available from the literature, where targets were examined with more than two IFNs. Kim et al. published the activity data of WT, Sniper, e-, evo-, Hypa- and SpCas9-HF1 on 6,481 target sequences that were suitable for our analyses^34^. These data confirmed the same activity/fidelity order of these five IFNs as we reported in this study. We analysed these data in silico from more than 32,000 (5 x 6,481) data points and found that these target sequences also tightly follow the cleavage rule (G-mean=0.981, Fig. 4c-e). This study also provided off-target cleavage data, of which we analysed the results from 30 sgRNAs on perfectly matching target sequences along with 1,800 off-target sequences containing all possible one-nucleotide mismatches for all nucleotide positions of the target and for all possible types of nucleotide change. The analyses confirmed our conclusion, that targets from different cleavability ranks require IFNs with correspondingly different fidelity ranks for specific cleavage (Fig. 4f). Here again, selecting the IFN that is closest to a target-matched one substantially increased the accuracy of IFN-editing (Fig. 4g). However, as seen in Figure 4g, there are still considerable off-target effects remaining when using only these 5 IFNs. This is consistent with the idea that a larger number IFNs can ensure a better resolution, and thus, provide an appropriate fit for more targets from the same set of target sequences. All the above results demonstrate that the sequence contributions of the targets in combination with the effect of the fidelity-increasing mutations of the IFNs primarily regulate on-target and off-target cleavage. These features appear to be universal; not specific to one cell-type or assay-type, and it applies to all variants and targets tested.

**Figure 3.**
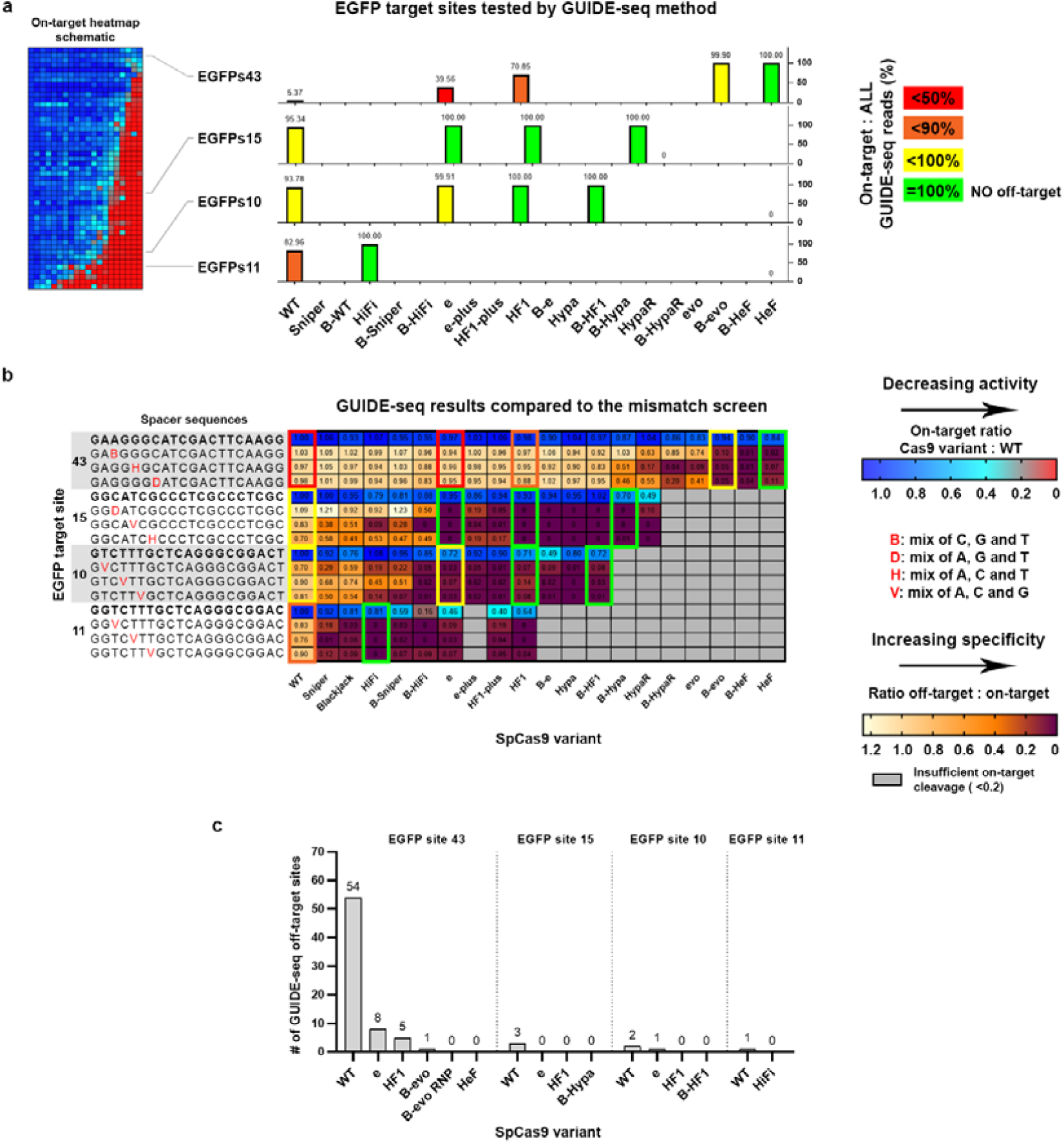
The genome-wide off-target effects of IFNs vary according to their mismatch tolerance, when compared on the same targets. **a-c,** Four targets and various IFNs were selected from the on-target and mismatch screen of Figure 2a and 2d and tested for genome-wide off-target events by GUIDE-seq. **a,** Overall on- target cleavage specificity of an IFN on a given target is expressed by the percentages of the on- target reads divided by all reads captured by GUIDE-seq (see calculations in Supplementary Table 6). Schematical heatmap of Figure 2a shows the position of the selected EGFP target sites within the ranking. **b,** Genome-wide off-target data fit appropriately into the results of the mismatch screen for the targets tested. Heatmap presented here is a segment from Figure 2d. The colour of the frames indicates the overall on-target specificity of an IFN on the given target as indicated in panel (**a**). **c,** Bar chart of the total number of off-target sites detected by GUIDE-seq for data shown in panels (**a-b**). For EGFP target 43, B-evo in ribonucleoprotein (RNP) form is also shown, which demonstrates that IFNs from higher fidelity ranks can be applied in RNP form to further increase specificity. **a-c,** Data is related to Figure 9, Supplementary Figures 5 and 7. Target sequences, raw, processed and heatmap disruption data, NGS and GUIDE-seq data are reported in Supplementary Tables 1-6.

**Figure 4.**
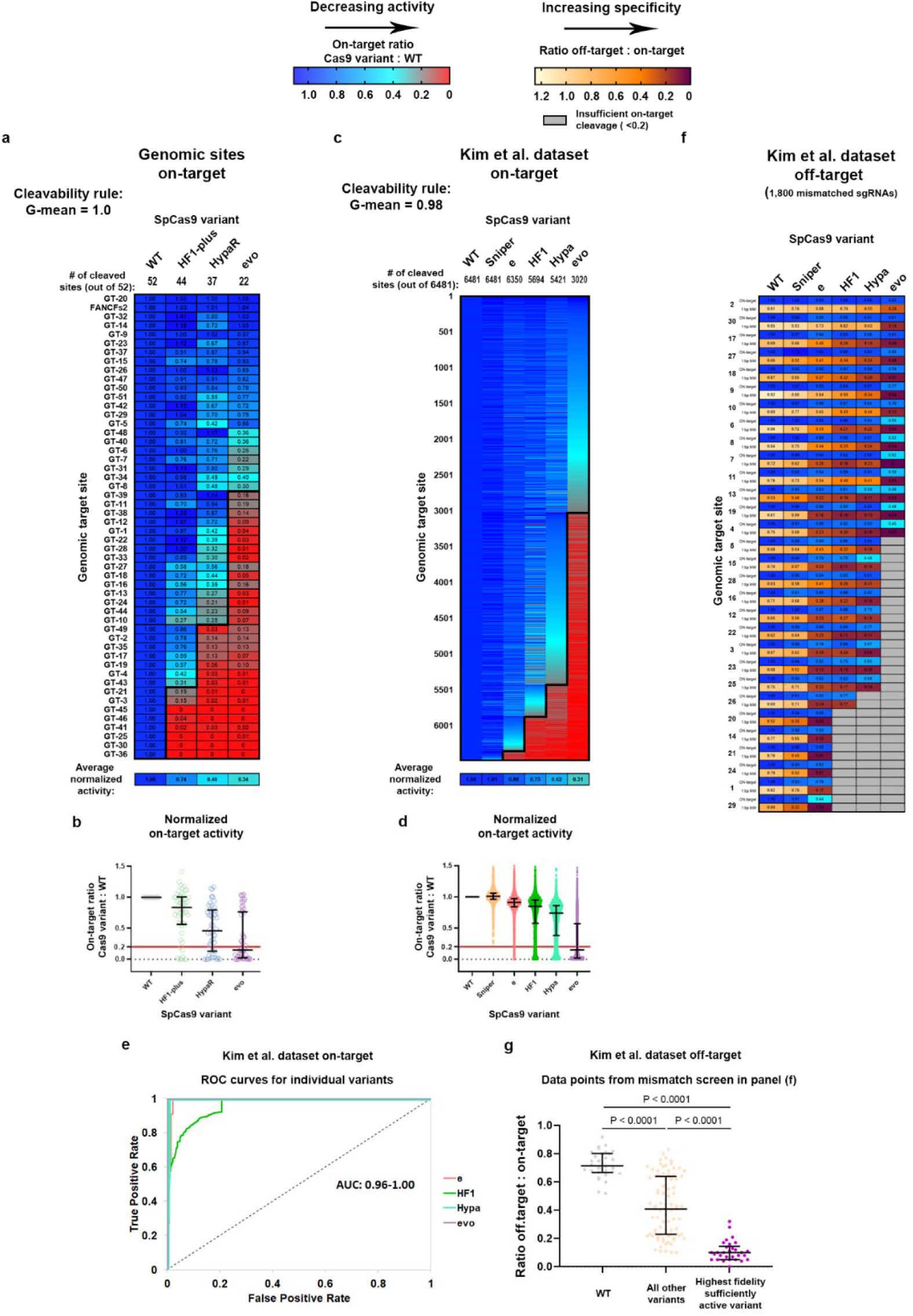
The cleavage rule still applies in a different cell line (HEK293) and on endogenous target sites examined by NGS. Heatmaps show the normalized value of the percentage of genome modification induced by SpCas9 variants with (**a, c**) perfectly matching and (**e**) partially mismatching 20G-sgRNAs. **a, c,** The bold line indicates the dividing line defined by the cleavage rule between the classes of cleaved and not-cleaved values. The G-mean value indicates how well the data points above and below the bold line correspond to the cleaved and not-cleaved (<0.20 activity normalized to WT) experimental values. **b, d,** Normalized on-target genome modification rates of various SpCas9 variants presented on a scatter dot plot. The sample points correspond to data presented in panel (**a**) and (**c**), respectively. Continuous red line indicates 0.20 normalized disruption activity, under which we consider the IFNs not to be active on a given target. The median and interquartile range are shown; data points are plotted as open circles representing the mean of biologically independent triplicates. Statistical significance was assessed by Friedman test and is shown in Supplementary Table 9. **c-d,** The data are compiled from experiments from Kim et al.^34^ and contain only selected sequences to avoid the effect of 5’ extended sgRNAs known to diminish the activity of all IFNs except Sniper and the SpCas9 variants containing the Blackjack mutations^21, 28^ (details can be found in Materials and Methods section and in Supplementary Table 7). The G-mean value of (**a**) 1.00 and (**c**) 0.98 (only 171 out of 32,405 data points are outliers) confirms that the cleavage rule is the main factor determining the activity of IFNs on genomic target sequences. **e,** The ROC curves verify that the order of the target sequences, determined by the cleavage rule, competently separates the classes of the cleaved and not-cleaved targets based on the normalized genome modification values of each individual variant from panel (**c)**. **f,** Mismatch screen of six nuclease variants with 30 sgRNAs on either perfectly matching or one-base mismatching target sequences. Data are compiled from experiments from Kim et al.^34^ and contain outcomes on 1,800 off-target sequences (details can be found in Materials and Methods section and in Supplementary Table 7). **g,** Matching IFNs and targets further increases the specificity of editing. The median and interquartile range of data points that are selected from panel (f) is presented as indicated; n=30, 87, 30, respectively. Dots are shown for each variant and target pair, where the on-target activity exceeded 70%. **a-g,** Target sequences, heatmap, NGS and data selected from Kim et al. and statistical details are reported in Supplementary Tables 1, 5, 7 and 9.

### The cleavage rule is discernible in the *in vitro* activities of IFNs

Next, we investigated whether the sequence contributions of the targets directly affect the cleavage activity of the variants, or they may derive solely from cellular effects. 21 targets from various cleavability ranks from Figure 2 were examined in an *in vitro* plasmid cleavage assay employing the purified ribonucleoprotein (RNP) complex of the WT SpCas9 and of either B-SpCas9-HF1, a variant from the middle of the fidelity ranking, or B-evoSpCas9 from the higher ranks (Fig. 5 and Supplementary Fig. 6). Figure 5c reveals that target sequences impact the activity of SpCas9s differently yielding a more than a magnitude difference in the cleavage rates in case of each variant, consistent with an earlier report^36^. Fidelity-increasing mutations decrease the activity of B-evoSpCas9 more than that of B-SpCas9-HF1 in a target-dependent manner. Most importantly, the combined effect of sequence contributions and fidelity-enhancing mutations is not only apparent *in cellulo*, but also *in vitro*, therefore it directly affects the cleavage activity of SpCas9s.

**Figure 5.**
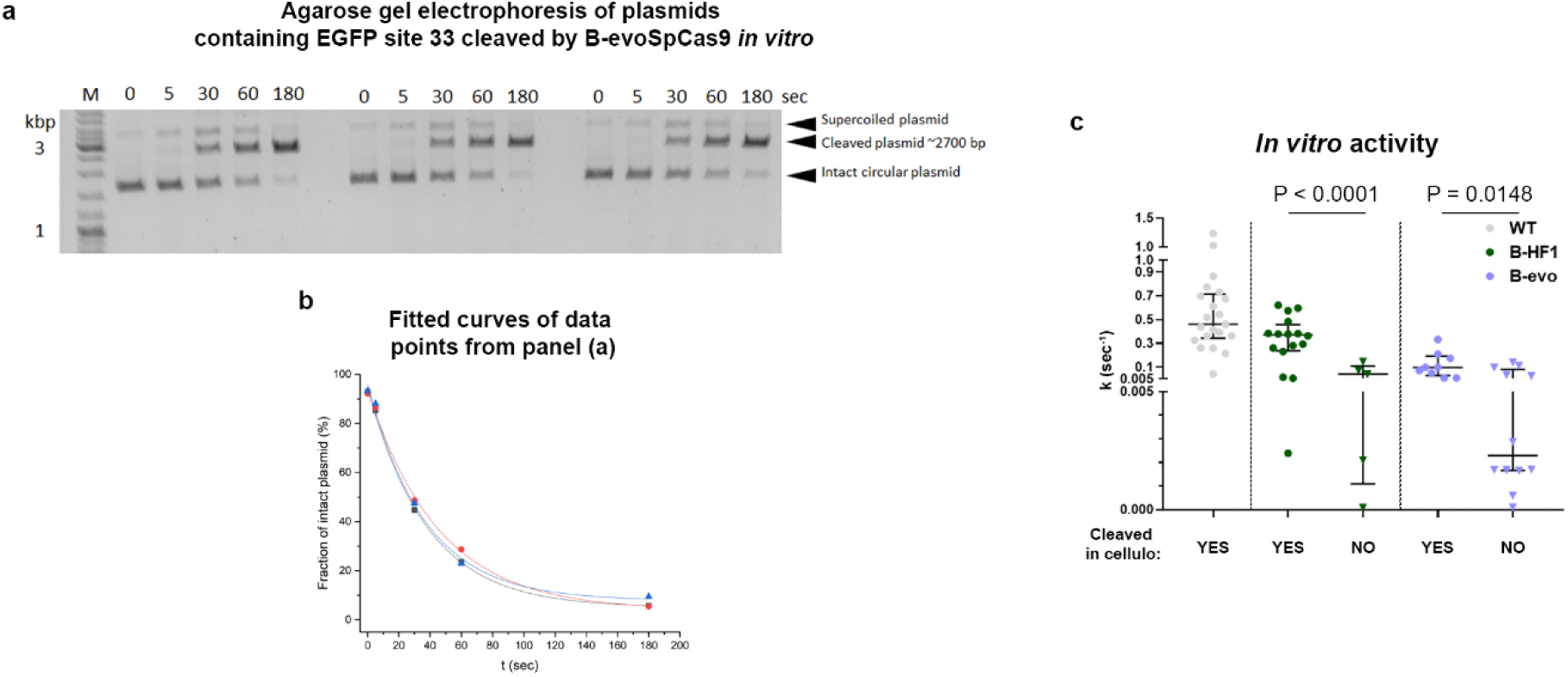
Sequence contributions are evident *in vitro*, supporting that it directly affects the enzymatic activity of IFNs. The effect of increased fidelity mutations and sequence contributions seen *in cellulo* also manifests in the rate constants of *in vitro* cleavage activities of the variants employing 21 targets of Figure 2. **a,** Representative agarose gel showing the activity of B-evoSpCas9 variant in a plasmid-cleaving *in vitro* assay at different timepoints in triplicates. **b,** Plot showing values representing the consumption of the intact circular plasmid (not-cleaved) derived from the intensity of bands from the representative agarose gel in panel (**a**). Exponential curves were fitted to the timepoints of each replicate separately. **c,** The average k values of three individual fits are shown. The median and interquartile range are shown; data points represent the mean of the fitted k value triplicates; n=21, 16, 5, 9, 12, respectively. **a-c,** Data related to Supplementary Figure 6. Target and primer sequences, *in vitro* data and statistical details are reported in Supplementary Tables 1, 8 and 9.

Two other arguments also support indirectly that the cleavage rule results from a direct interaction between IFNs and targets. (i) The EGFP disruption experiments demonstrate that the observed differences in the cleavability of the targets by IFNs (but not WT) *in cellulo* does not result from the location of the targets (whether in chromatin, coding or non-coding regions) or from the transcript levels when they are in a transcribed region, since the targets shown in Figure 2a are all located within the EGFP sequence integrated into one location of the genome. (ii) It can also be argued that higher-ranking IFNs may have WT-like activity at high-ranking targets, because cleavage at these targets is saturated, so their reduced activity is not apparent. This scenario can be probed by titrating such targets with plasmids that express higher-ranking IFNs by transfecting different amounts of plasmids. In fact, we carried out this experiment in our recently published study (bioRxiv: https://doi.org/10.1101/2022.05.27.493683) where Supplementary Figure 1e shows that the activity of the highest fidelity IFNs decreases in parallel with WT and lower-ranking IFNs, which argues against a saturating effect.

Taken together, these data suggest that the combined direct effect of target sequence contributions, fidelity-increasing mutations and mismatches on SpCas9 activity result in the emergence of the cleavage rule. When the effect of sequence contributions is much larger than the effect of fidelity-increasing mutations, then not only target cleavage occurs with substantial off-target effects, but also the impact of other intrinsic and cellular factors is more pronounced, modulating the level of WT-normalized activity, typically between 70% and 120% (Fig. 7b and 8).

### SpCas9-NG and xCas9 do not obey the cleavage rule

With the established knowledge that an appropriate set of IFNs rather than any individual variant is necessary for reaching maximal specificity universally for any target, it would be a particularly useful idea to create an alternative set of IFNs with altered-PAM specificities. This would increase the accessibility of target sequences by the recognition of targets with an NG-like PAM sequence instead of the canonical NGG. Such variants, like SpCas9-NG and xCas9, have also been reported to possess increased fidelity and relatively low activity^37, 38^. In order to create IFNs that belong to the lower fidelity ranks but with NG PAM specificity, the activity of the SpCas9-NG or xCas9 would need to be increased. Some mutations in xCas9 have been hypothesized to primarily increase the fidelity of the variant, instead of contributing to the altering of the PAM specificity^39^. Our efforts to increase its activity by replacing these mutations with the wild type amino acids were unsuccessful (Fig. 6a). As an alternative solution, we applied fidelity decreasing mutations^22^ and demonstrated their effects on the activity of HypaR- SpCas9, an IFN with activity close to that of SpCas9-NG and xCas9, and on targets that are in the cleavability ranks just on the border of cleaved or not cleaved by HypaR. However, when we introduced them to SpCas9-NG or xCas9, the mutations did not decrease their target selectivity on the tested sequences (Fig. 6b- c). Intriguingly, SpCas9-NG and xCas9 do not fit in with the pattern formed by the rest of the IFNs (Fig. 6d). They do not strictly obey the cleavage rule of the targets (Fig. 6e). In this respect, it would be interesting to see other PAM-altered variants, such as ones developed by Kleinstiver and co-workers, whether they also behave like SpCas9-NG and xCas9^40–45^. These results also highlight, that a variant with reduced activity, even with seemingly increased specificity, does not automatically qualify for the IFN ranking, and that the cleavage rule resulting the pattern seen in Figure 2 is not something self-evident.

**Figure 6.**
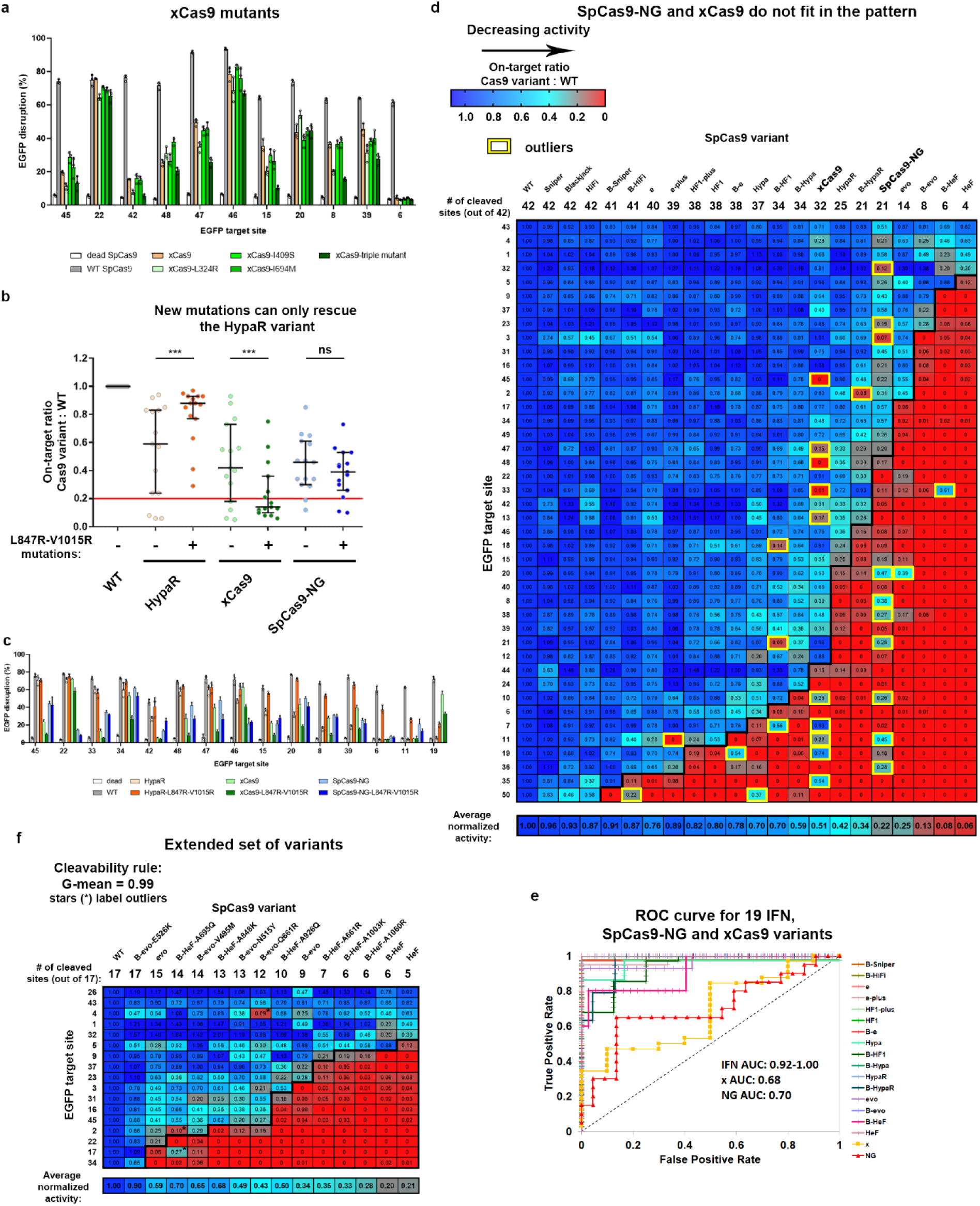
New increased fidelity SpCas9 variants in the higher fidelity ranks with in-between activity/fidelity a, c, The results of an on-target EGFP disruption assay for WT and different SpCas9 variants on various target sites shown on a column graph. Means and SD are shown; n=3 biologically independent samples (overlaid as white circles). **a,** Reverting three mutations of xCas9 back to the WT residues, suggested by Guo et al.^39^ as being responsible for its increased fidelity and target-selectivity, did not increase its activity the way we expected. Statistical significance was assessed by RM one-way ANOVA and shown in Supplementary Table 9. **b,** Normalized on-target activities of various SpCas9 variants presented on a scatter dot plot. The sample points correspond to data presented in panel (**c**). Continuous red line indicates 0.20 normalized disruption activity, under which we consider the IFNs not to be active on a given target. The median and interquartile range are shown; data points are plotted as open circles representing the mean of biologically independent triplicates. **d, f,** Heatmaps show the normalized EGFP disruption activities of SpCas9 nucleases with perfectly matching 20G-sgRNAs. The bold line indicates the dividing line defined by the cleavage rule between the classes of cleaved and not-cleaved values. **d,** SpCas9-NG and xCas9 do not strictly obey the cleavage rule when fitted on the heatmap of Figure 2a (only 42 EGFP target sites were tested here), even though some of the targets for which the order were not determined by the 19 IFNs were reordered (compared to Fig. 2a heatmap) to favour the accommodation of SpCas9-NG and xCas9 into the cleavage map. These results explain the failure to develop an IFN series with a looser PAM requirement and emphasize that the cleavage rule identified here is non-trivial and does not apply to all other SpCas9 variants with reduced activity or increased fidelity. **e,** The ROC curves demonstrate that the order of the target sequences, determined by the cleavage rule, competently separates the classes of the cleaved and not-cleaved targets in case of the IFN variants, but xCas9 (AUC: 0.68) and SpCas9-NG (AUC:0.70) do not appear to strictly follow the cleavage rule, emphasizing that the rule is not self-evident. **f,** New IFNs provide a finer resolution within the higher fidelity region of the IFN ranking between evo- and HeFSpCas9, and their activities on these targets also strictly follow the cleavage rule. For these experiments, we selected targets from the higher cleavability region of the target ranking in Figure 2, as these were expected to be the point of distinction between the new variants, and therefore facilitate the ordering of these IFNs and their fitting into the already existing ranking. This is the reason for these high fidelity IFNs showing much higher normalized disruption activities, than they would on randomly selected targets. **a-f,** Target sequences, raw, processed, heatmap disruption data and statistical details are reported in Supplementary Tables 1-4, 9.

From here, we progressed parallel, on the one hand, (i) with the generation of more IFNs with intermediate fidelity for ranks with lower resolution, while on the other hand, (ii) proceeding with this set of 19 variants to assess if this panel was large enough to demonstrate that target-matched IFNs facilitate genome editing with maximal specificity i.e., without any detectable genome-wide off-target.

### Extending the set of IFNs with ten new variants with intermediate fidelity

One of the main conclusions of our study is that a full series of IFNs is needed in order to be able to provide highly specific editing in general, for any given target regardless of its cleavability rank. However, the distribution of our set of 19 increased fidelity SpCas9s is not spread evenly across the full range of the fidelity ranking. There are more IFNs in the lower/medium fidelity range and some of them do not or just marginally differ in on-target activity/fidelity. In contrast, there are only a few options for targets requiring nucleases with higher fidelity. Therefore, to provide a better resolution of the available IFNs in these higher fidelity ranks, we reverted several single mutations^22^ in B-evo- and B-HeFSpCas9 to the original WT amino acids creating ten new variants with the intended intermediate fidelity and target-selectivity (Fig. 6f). These variants provide additional tools for editing those targets from the high cleavability ranks where the panel of the 19 variants may not provide a sufficiently matching IFN.

### Identifying the target-matched variants

Finally, the most important result of this study is that by employing target-matched IFNs we are able to ensure maximal specificity editing for practically any target sequence, that is accessible to WT SpCas9, without any genome-wide off-target effects. Several clever and effective genome-wide off-target detecting methods have been developed *in vitro* and *in cellulo*^11, 20, 46–58^. While *in vitro* methods tend to report more off-target sites, they are prone to identifying a high number hits with uncertain relevance and require extensive validations. In contrast, GUIDE-seq, likely the most widely used approach, is reported to have the highest validation rate amongst genome-wide methods^49^ and its sensitivity is comparable to or even higher than that of amplicon sequencing (NGS) (Supplementary Fig. 7a)^59^. Thus, given the rather large number of pairs of target and variant to be tested, in this study we have relied on GUIDE-seq to monitor the off-target activity of the nucleases. To identify the target-matched variants for a given target and to select the optimal one for the maximal specificity, we used a two-step approach by exploiting the observed cleavage rule of the targets (Fig. 7a). In the first step, we measure the on-target activity of WT and three IFNs (e-plus, B-Hypa and B- HypaR), that divide the target range in Figure 2a into four proportional sections based on the fraction of the targets they can cleave, to identify which one has the highest fidelity, that still shows sufficient efficiency. In the second step, a few additional IFNs, neighbouring the one selected in the first step, are tested for on-target activity to identify target-matched variants. Finally, out of these variants, the optimal target-matched variant with the maximum specificity is selected and/or confirmed by GUIDE-seq. We show this strategy on HEK site 1, 2, 3 and *VEGFA* site 1, that have been analysed by GUIDE-seq previously^20^. Figure 7b demonstrates that with the panel of 19 IFNs, all four targets could be edited without any genome-wide off-target effect detected by GUIDE-seq.

**Figure 7.**
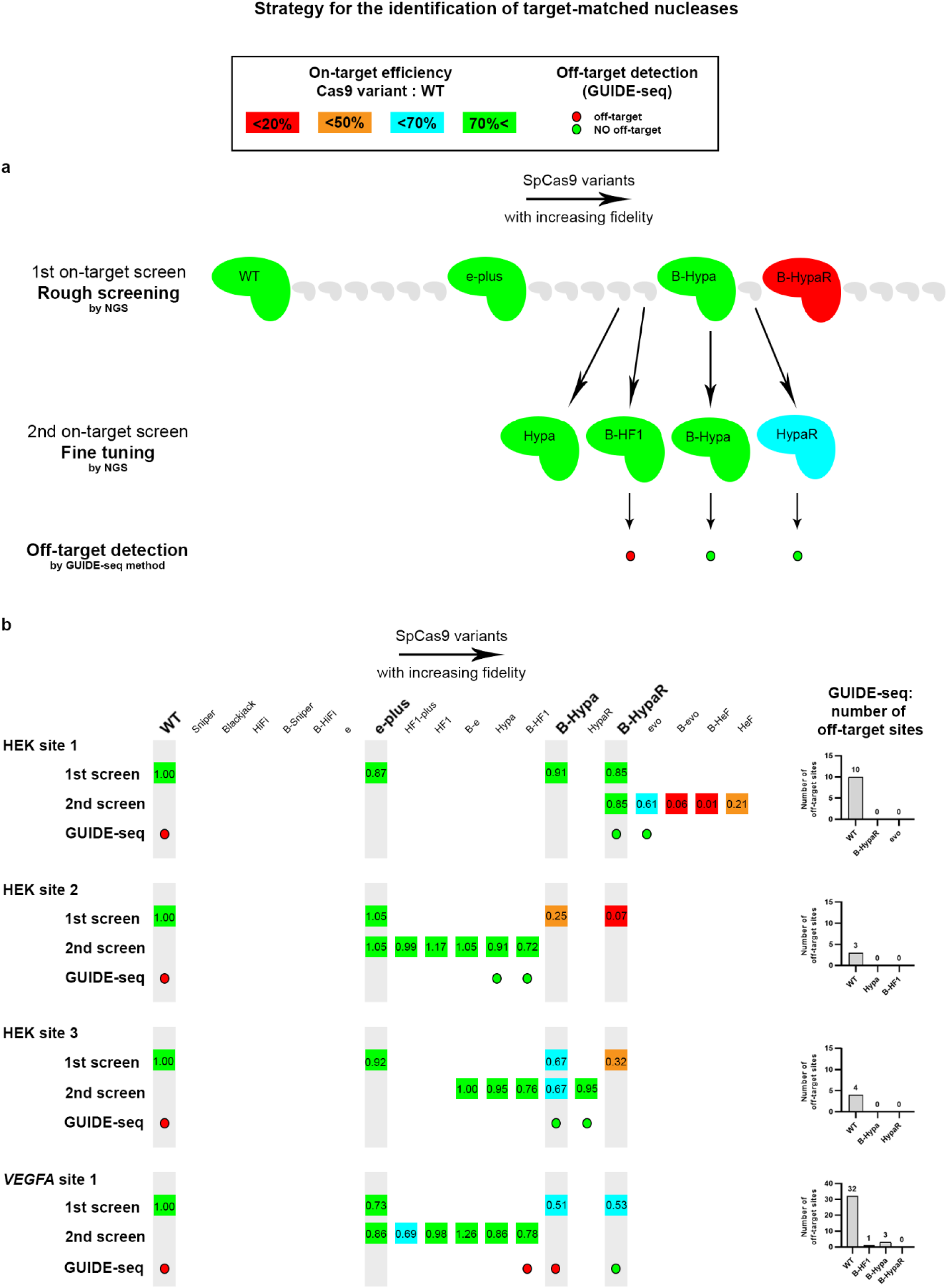
The optimal target-matched SpCas9 nuclease showing efficient on-target editing and no off-target effects is identified for each target using a two-step approach a, Schematic representation of the two-step strategy used. In the 1^st^, rough on-target screen, the WT and three IFNs are used, which divide the target ranking range into approximately four proportional sections. The 2^nd^, fine-tuning on-target screen identifies the target-matched variants; for this, we used at least four variants that are positioned right next to the IFNs with the highest fidelity identified in the rough screening that are still sufficiently active. If necessary, two-to-three sufficiently active (normalized activity above 50-70% depending on the application under consideration) target-matched variants can be screened for the absence of genome-wide off-targets. **b,** The identification of the target-matched variants that provide appropriate editing without any genome-wide off-target is demonstrated on four targets that had been tested in Tsai et al.^20^. The numbers in the coloured squares indicate the value of the percentage of on-target genome modification normalized to WT. The total number of off-target sites detected by GUIDE-seq are shown for each target in bar charts on the right side of the panel. Data related to Figure 9, Supplementary Figures 8 and 9. Target sequences, NGS and GUIDE-seq data are reported in Supplementary Tables 1, 5 and 6.

### Editing challenging targets efficiently without any detectable off-targets

We have already shown the usefulness of the application of target-matched IFNs on 4 EGFP targets and then on another 4 genomic targets (Fig. 3 and 7b), however, in order to draw meaningful conclusions, instead of simply adding extra arbitrarily selected targets, we challenged our approach by examining all problematic target sequences reported in the literature that have been failed to be edited by any of the IFNs without genome-wide off-targets (Supplementary Table 6: Data from other studies). Most studies characterising IFNs focused on the same or an overlapping set of targets in order to provide the new variant with a relevant comparison to the preceding ones. Thus, these studies together ended up examining the same targets with a number of variants and by chance some of these tests involved a target-matched variant for several of the targets. In some cases, the target could not be edited without off-targets in spite of all efforts, simply because the existing/tested variant IFN set did not contain the target-matched variants. Here we tested our approach on the seven targets on which the former IFN-studies failed to provide efficient and off-target-free editing^21, 23, 24, 27, 29^. Since prior information exist for these targets, in case of some of the targets we performed only one round of on-target screen to identify target-matched nucleases. In contrast to previous studies, using the understanding of the cleavage rule, the extended set of IFNs developed here and by the straightforward identification of the target-matched variants, we managed to successfully edit all seven challenging targets without any detectable genome-wide off-target (Fig. 8 and Supplementary Fig. 7b-e). Most impressively, *VEGFA* site 2, 3 and *FANCF* site 2, which have been previously failed by 7, 4 and 7 IFNs, respectively^21, 23, 24, 27^, were also edited without genome-wide off-targets by using target-matched nucleases in RNP form. These results project that by the use of an appropriate set of IFNs, such as the one we generated, virtually any target from any rank can be edited with greatly enhanced specificity, without any off-target effect (experiments summarized in Figure 9 and detailed in Figures 3, 7, 8 and 10, Supplementary Figures 5, 7, 8 and 9). The greatest benefit of these results is likely to be realised in therapeutic applications of genetic engineering, where maximum specificity and safety are required.

**Figure 8.**
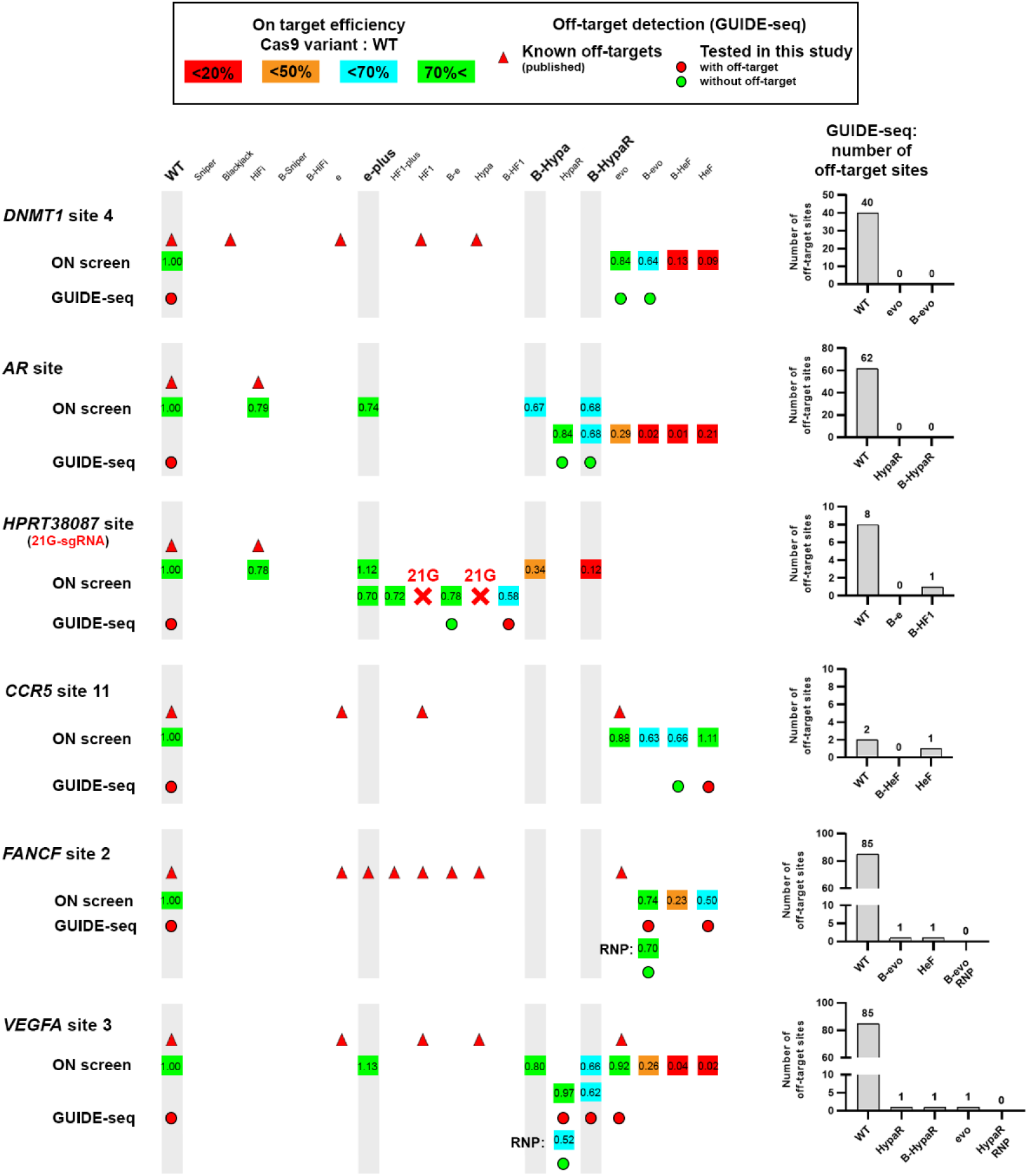
Even repetitive, non-typical sequences can be edited without off-targets by employing optimal target-matched IFNs. Targets shown here had previously only been edited by IFNs with off-targets, red triangles indicate these previous attempts. Here, they were all successfully edited without any genome- wide off-targets detectable by GUIDE-seq using target-matched IFNs. The numbers in the coloured squares indicate the value of the percentage of on-target genome modification normalized to WT (measured by NGS). GUIDE-seq was performed with two to three target- matched IFNs that reached at least 50% normalized on-target editing. Some targets can be edited with no detectable genome-wide off-targets by more than one target-matched IFN. We used a two-step approach to identify potential IFNs that could be used to edit a given target without any off-target, except for three targets, for which we had sufficient information to skip the rough screen; the first step of the approach. In the case of *FANCF* site 2 and *VEGFA* site 3 a lower fidelity variant in RNP form showed higher specificity than a higher-ranked variant that was expressed from a plasmid. Bar chart of the total number of off-target sites detected by GUIDE-seq are shown on the right side of the panel. Data related to Figure 9, Supplementary Figures 8 and 9. Target sequences, NGS and GUIDE-seq data are reported in Supplementary Tables 1, 5 and 6.

**Figure 9.**
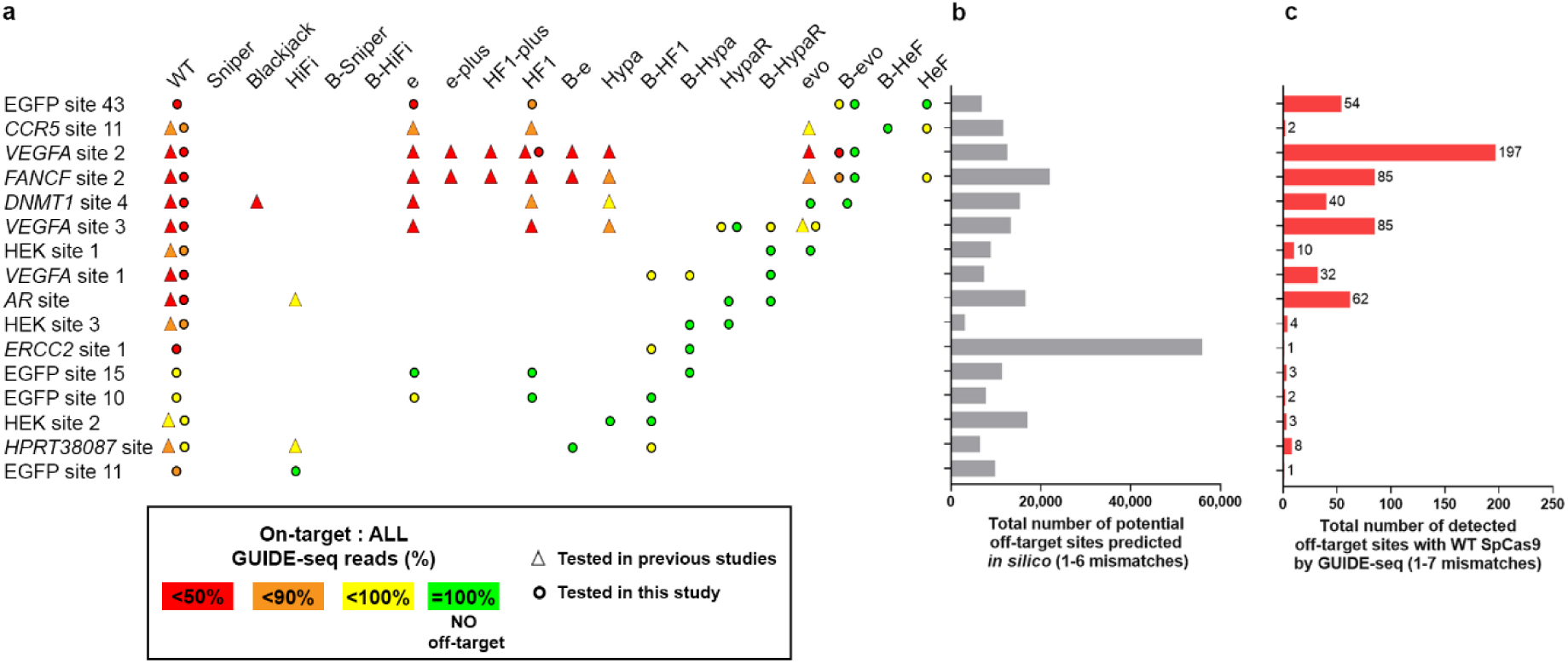
Target-matched nucleases show high efficiency without any genome-wide off-target for all targets tested in this study regardless their ranking. Summary of targets edited by IFNs, examined in this study. For 16 target sites, including all those challenging targets where previous attempts with IFNs had failed, we were able to perform editing without any off-target detectable by GUIDE-seq. The GUIDE-seq results are shown from experiments of both previous studies (filled triangles) and our studies (filled circles) for the selected individual variants on the indicated targets. Overall specificity on a given target is indicated by different colours, as stated on the figure. When a target is edited with the same variant in more than one literature study, then the one with the highest overall GUIDE-seq read is displayed. The ranking of the targets and either **b**, the number of their predicted off-target sites or **c**, the detected off-target sites using the WT SpCas9 are weakly related (for details see Supplementary Table 10). Data are related to Figures 3, 7, 8 and 10, Supplementary Figures 5, 8 and 9.

### Correcting a clinically relevant mutation without any detectable off-target

We also attempted to correct a clinically relevant mutation in a patient-derived cell line to present the power of the method on a relevant target site that we had no prior knowledge of. Cells with a defective mutation were derived from a patient with Xeroderma pigmentosum, a rare genetic disorder without any cure to date^60^. They harbour a C>T substitution, which results in the change of Arg-683 to Trp disrupting the function of the *ERCC2* gene. Patients with Xeroderma pigmentosum are extremely sensitive to the ultraviolet range of sunlight as a result of dysfunctional DNA repair, which often leads to the development of skin cancer and early death at a young age^61^. We located the target sequence nearest to the mutation, identified the optimal target-matched IFN and corrected the mutation with B-HypaSpCas9 in 10.7% of the cells using single-stranded DNA oligonucleotides without any detectable off-target effect, indicating the high potential of our approach (Fig. 10).

**Figure 10.**
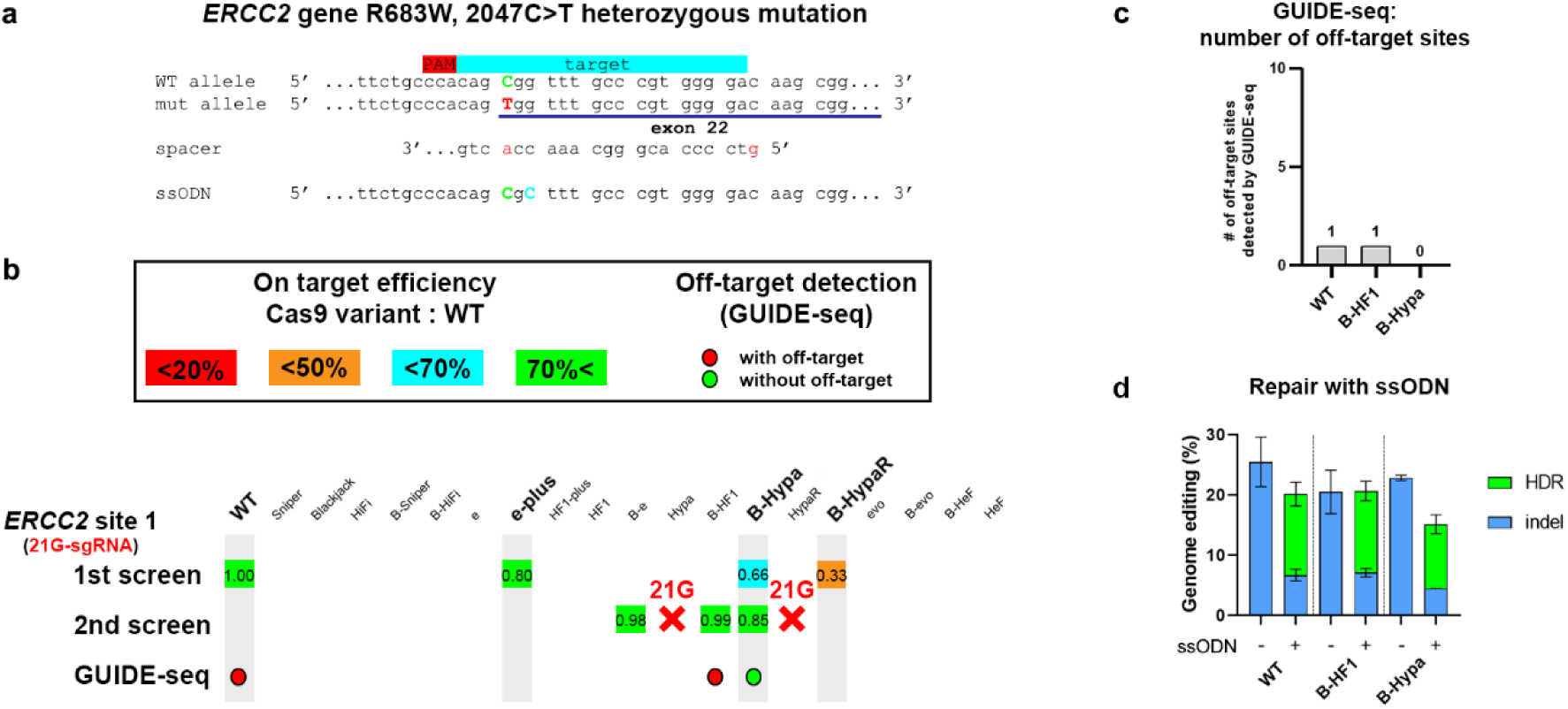
Correcting a clinically relevant mutation without off-target cleavage using the two-step approach used earlier a, The strategy to correct the mutation causing Xeroderma pigmentosum in patient-derived fibroblast cells is shown, including the sequence environment of the mutation (disease-causing mutation – red letter, WT nucleotide– green letter, silent mutation – blue letter). **b,** By using the two-step approach we identified B-HypaSpCas9 for editing without any detectable genome-wide off-target. Values in the coloured squares show the normalized value (to WT) of the percentage of on-target genome modifications measured by NGS. Hypa and Hypa-R being non-Blackjack- IFN exhibited diminished (<0.2) activity with the 21G-sgRNA. **c,** Bar chart are shown of the total number of off-target sites detected by GUIDE-seq. **d,** B-HypaSpCas9 with single strand oligo nucleotide repair using HDR enhancer M3814^62^ provides WT-like level of correction of the R683W (2047C>T) mutation. Means and SD are shown; n=3. **a-d,** Data related to Figure 9, Supplementary Figures 8 and 9. Target sequences, NGS and GUIDE-seq data are reported in Supplementary Tables 1, 5 and 6.

## Discussion

There are two major achievements in this study; on the one hand we recognised the cleavability ranking of the targets and established the cleavage rule, that governs the outcome of the interactions between IFNs and targets, and then, used this knowledge to develop new IFNs that fill the gaps in the fidelity ranking of the variants so that we can provide a suitable IFN for targets from any cleavability ranking. On the other hand, by exploiting the cleavage rule and the newly developed set of IFNs we demonstrated that both maximal specificity (i.e., no detectable genome-wide off-targets, as assessed by GUIDE-seq) and efficient cleavage can be expected to be achieved universally for any target. We note three issues related to the cleavage rule; (i) The cleavage rule is the combined result of the impact of the fidelity-increasing protein mutations in SpCas9 variants and of the actual sequence of the targets/spacers on the activity of SpCas9. Although the cleavage rule was extremely strictly followed by the IFNs in every dataset examined (G-means range between 0.98 and 1.0), other factors also influence the activity of the variants resulting in some outliers with unexpected on-target cleavage activities or residual off-target effects. Nevertheless, the fact that the effect of fidelity-increasing mutations and target sequence contributions is so clearly observed, argues that these are the main factors. (ii) The cleavage rule perfectly separates IFNs working and non-working on a target, but this does not mean that working IFNs necessarily show continuously decreasing normalized percentages according to their position in the ranking. These values are typically above 70-80%, although lower values occasionally occur. (iii) The cleavage rule predicts that maximum specificity is achieved with the IFN having the highest fidelity and sufficient activity on a given target. Since other factors may modify this scenario, when maximum fidelity is required it is recommended that the two IFNs with the highest fidelity should be tested. Indeed, when no RNP form was used for maximum specificity editing of the target, we found only two cases in the whole study (*HPRT38087* and *CCR5*s11) where the highest-ranking IFN did not show maximum specificity editing (i.e.: without genome-wide off-targets). IFNs in the RNP form having higher specificity than they have when they are introduced into the cells as a plasmid, do not fit the same position of fidelity ranking as in the case of plasmid-based transfection. For simplicity, we placed them to the position of the corresponding IFN used in plasmid form. (iv) The recognition of the cleavability ranking of SpCas9 targets may inspire researchers to revisit some structural and mechanistical studies of SpCas9 that are typically performed on a single target, by examining targets of different ranks to cross-check their conclusions. (v) Furthermore, *in vitro* data showed that sequence contributions resulting in the cleavability ranking of the targets directly affect the cleavage activity of these SpCas9 nuclease variants, however, further research is required to understand what sequence features exactly affect the cleavage of the IFNs.

Regarding of the use of target-matched IFNs, we highlight the following. (i) By employing two on-target screening steps, the nucleases with the highest fidelity rank and with sufficient activity for the given application can be chosen without having to resort to any genome-wide off-target assessing method. These variants provide highly specific and efficient editing. However, when maximal specificity is required, it may be more prudent to examine the two best candidate variants, as several factors affect the activity of an IFN, resulting in unexpected outliers with residual off-targets in some cases. (ii) Here, we showed that practically any target that is efficiently edited by the WT SpCas9 can be expected to be edited efficiently, without off-targets by employing target-matched IFNs, thus considerably increasing the potential of genome engineering in terms of safety and efficiency when high specificity is required, such as gene therapeutics. (iii) Although the majority of the off-target mutations may have no detrimental consequences, the few that do, still uphold substantial threat as *ex vivo* and *in vivo* therapeutic applications involve millions to billions of cells. The routine use of gene therapy further increases the risk by thousands of folds, in contrast to a single treatment. (iv) Furthermore, the off-target cleavages by the nuclease even in innocuous positions can still pose a significant risk, as double-strand breaks at off-target positions increase the chance of chromosomal translocations that can also lead to cancerous transformation^13, 63^. (v) For safe therapeutic procedure the aim needs to be maximal specificity, possibly beyond the about 0.1% detection limits of current methods^49, 59^ for the assessment of off-targets. Since a target may be edited without detectable off-targets by multiple IFNs, in such cases, as a general practice, the target-matched IFN with the highest fidelity should be identified and applied. Furthermore, target-matched IFNs are compatible with many other approaches (e.g., RNP form or dRNA^64^) for decreasing off-targets, and it is worth considering their application to maximize specificity even in cases that would fall under the detection limits of off-target detecting methods. (vi) The use of target-matched IFNs may also be beneficial in base and prime editing^65–67^. These methods work with substantially less Cas9 dependent off-targets than nucleases, nevertheless, they also rely on cleavage, i.e., the nickase activity of SpCas9. The nickase versions of IFNs seem to exhibit the same sensitivity to the sequence contributions of the targets^32, 68^, thus applying target-matched IFN base and prime editors may decrease off-target editing of current editors to a non-detectable level and further. (vii) The identification of target-matched IFNs for a given target has proved to be straightforward here, still, a predictive algorithm, which could identify the target-matched IFNs for specific targets could further simplify this process and make it less labour intensive. Unfortunately, prediction programs to date are not accurate enough to suggest a reliable choice. Large cleavage activity data for a considerable number of IFNs from all fidelity ranges of the ranking should be generated for the development of an appropriate prediction tool. (viii) CRISPR-based gene therapy has reached the stage of clinical trials for some clinically tractable monogenic disorders^1–3^. The editing of a specific target can reach clinical applications after extensive and very expensive optimisation including aspects of efficacy and specificity. However, a considerable fraction of mendelian disorders is N=1 diseases^69^ where only one patient carries a given mutation, making the high cost of extensive optimisation prohibitive. While several issues need to be solved^69^, our methodology also seems particularly promising for such N=1 diseases, where we offer an effective way for the optimisation of critical steps of the procedure of gene therapy resulting in acceptable safety in a short timeframe with affordable cost, contributing to the democratisation of the CRISPR technology.

In conclusion, the translation of advances in CRISPR technology into clinical applications faces several challenges in terms of the efficiency of the modification, the delivery of the tools *in vivo* as well as various undesired, non-intended modifications affecting the genome. Our approach diminishes one of these obstacles, the appearance of off-target edits, and therefore it provides an exceptionally high precision tool for research and therapeutic applications.

## Materials and methods

### Materials

Restriction enzymes, T4 ligase, Dulbecco’s modified Eagle Medium DMEM (Gibco), fetal bovine serum (Gibco), Turbofect, TranscriptAid T7 High Yield Transcription Kit, Qubit dsDNA HS Assay Kit, Taq DNA polymerase (recombinant), Platinum Taq DNA polymerase,0.45 µm sterile filters and penicillin/streptomycin were purchased from Thermo Fischer Scientific, protease inhibitor cocktail was purchased from Roche Diagnostics. DNA oligonucleotides, trimethoprim (TMP), chloroquine, polybrene, puromycin, calcium-phosphate and GenElute HP Plasmid Miniprep kit were acquired from Sigma-Aldrich. ZymoPure Plasmid Midiprep kit and RNA Clean & Concentrator kit were purchased from Zymo Research. NEBuilder HiFi DNA Assembly Master Mix and Q5 High-Fidelity DNA Polymerase were obtained from New England Biolabs Inc. NucleoSpin Gel and PCR Clean-up kit was purchased from Macherey-Nagel. 2 mm electroporation cuvettes were acquired from Cell Projects Ltd, SF Cell Line 4D-Nucleofector X Kit S were purchased from Lonza, Bioruptor 0.5 ml Microtubes for DNA Shearing from Diagenode. Agencourt AMPure XP beads were purchased from Beckman Coulter. T4 DNA ligase (for GUIDE-seq) and end-repair mix were acquired from Enzymatics. KAPA universal qPCR Master Mix was purchased from KAPA Biosystems.

### Plasmid construction

Vectors were constructed using standard molecular biology techniques including the one-pot cloning method^70^, E. coli DH5α-mediated DNA assembly method^71^, NEBuilder HiFi DNA Assembly and Body Double cloning method^72^. All SpCas9 variants were codon optimized the same way. Plasmids were transformed into NEB Stable competent cells or DH5alpha. For detailed cloning and sequence information see Supplementary Information. A list of sgRNA target sites, mismatching sgRNA sequences and plasmid constructs used in this study are available in Supplementary Table 1. The sequences of all plasmid constructs were confirmed by Sanger sequencing (Microsynth AG).

Plasmids acquired from the non-profit plasmid distribution service Addgene (http://www.addgene.org/) are the following:

pX330-U6-Chimeric_BB-CBh-hSpCas9 (Addgene #42230)^6^, eSpCas9(1.1) (Addgene # 71814)^22^, VP12 (Addgene #72247)^23^, sgRNA(MS2) cloning backbone (Plasmid #61424)^73^, pMJ806 (#39312)^7^, pBMN DHFR(DD)-YFP (#29325)^74^ and p3s-Sniper-Cas9 (#113912)^28^.

pX330-SpCas9-NG (#117919) was a kind gift from Hiroshi Nishimasu.

Plasmids developed by us in this study and deposited at Addgene are the following:

B-Sniper SpCas9 (#), B-HiFi SpCas9 (#) HypaR-SpCas9 (Addgene #126757), B-HypaR-SpCas9 (Addgene #126764).

### *In vitro* transcription

sgRNAs were transcribed *in vitro* using TranscriptAid T7 High Yield Transcription Kit and PCR-generated double-stranded DNA templates carrying a T7 promoter sequence. PCR primers used for the preparation of the DNA templates are listed in Supplementary Table 1. sgRNAs were purified with the RNA Clean & Concentrator kit and reannealed (95 °C for 5 min, ramp to 25 °C at 0.3 °C/s). sgRNAs were quality checked using 10% denaturing polyacrylamide gels and ethidium bromide staining.

### Protein purification

All SpCas9 variants were subcloned from pMJ806 (Addgene #39312)^7^ [except pET-HypaR- SpCas9-NLS-6xHis, which was subcloned in pET-Cas9-NLS-6xHis (Addgene #62933) plasmid]. For detailed cloning information and sequence information see Methods: Plasmid construction section, Supplementary Table 1 and Supplementary Information. The resulting fusion constructs contained an N-terminal hexahistidine (His6), a Maltose binding protein (MBP) tag and a Tobacco etch virus (TEV) protease site (except pET-HypaR-SpCas9-NLS-6xHis).

The expression constructs of the SpCas9 variants were transformed into *E. coli* BL21 Rosetta 2 (DE3) cells, grown in Luria-Bertani (LB) medium at 37 °C for 16 h. 10 ml from this culture was inoculated into 1 l of growth media (12 g/l Tripton, 24 g/l Yeast, 10 g/l NaCl, 883 mg/l NaH_2_PO_4_ H_2_O, 4.77 g/l Na_2_HPO_4_, pH 7.5) and cells were grown at 37 °C to a final cell density of 0.6 OD600, and then were cooled to 18 °C. The protein was expressed at 18 °C for 16 h following induction with 0.2 mM IPTG. Proteins were purified by a combination of chromatographic steps by NGC Scout Medium-Pressure Chromatography Systems (Bio-Rad). The bacterial cells were centrifuged at 6,000 rcf for 15 min at 4 °C. The cells were resuspended in 30 ml of Lysis Buffer (40 mM Tris pH 8.0, 500 mM NaCl, 20 mM imidazole, 1 mM TCEP) supplemented with Protease Inhibitor Cocktail (1 tablet/30 ml; complete, EDTA-free, Roche) and sonicated on ice. Lysate was cleared by centrifugation at 48,000 rcf for 40 min at 4 °C. Clarified lysate was bound to a 5 ml Mini Nuvia IMAC Ni-Charged column (Bio-Rad). The resin was washed extensively with a solution of 40 mM Tris pH 8.0, 500 mM NaCl, 20 mM imidazole, and the bound proteins were eluted by a solution of 40 mM Tris pH 8.0, 250 mM imidazole, 150 mM NaCl, 1 mM TCEP. 10% glycerol was added to the eluted sample and the His6-MBP fusion proteins were cleaved by TEV protease (3 h at 25 °C) (except pET-HypaR-SpCas9-NLS-6xHis). The volume of the protein solution was made up to 100 ml with buffer (20 mM HEPES pH 7.5, 100 mM KCl, 1 mM DTT). Proteins were purified on a 5 ml HiTrap SP HP cation exchange column (GE Healthcare) and eluted with 1 M KCl, 20 mM HEPES pH 7.5, 1 mM DTT. They were then further purified by size exclusion chromatography on a Superdex 200 10/300 GL column (GE Healthcare) in 20 mM HEPES pH 7.5, 200 mM KCl, 1 mM DTT and 10% glycerol. The eluted protein was confirmed by SDS-PAGE and Coomassie brilliant blue R-250 staining, and they were stored at -20 °C.

### Determining active SpCas9 quantity in solution

The quantification method was based on Liu et al.^75^. The quantity of active SpCas9 protein in solution was determined using EGFP target site 32, that has shown high cleavage activity with all three proteins tested based on previous experiments. The measurement procedure is as follows: The target plasmid was incubated for an hour with protein-sgRNA complex, in different concentrations. Concentrations were determined by spectrophotometry (Nanodrop OneC), and then the target site containing the plasmid (10 nM) and the SpCas9 protein were mixed in a ratio between 1:0.5 and 1:10, while the quantity of the sgRNA was twice that of the protein in each case. To terminate cleavage reaction, the inactivation solution (final concentration: 0.2% SDS, 50 mM EDTA) was added to the reaction mix at 80 °C. Samples were ran on a 0.8% agarose gel. Following densitometry (GelQuantNET, BiochemLabSolutions.com), the ratio of intact plasmid and total DNA was calculated for each sample. These values were plotted and fitted on a ‘One-phase exponential decay function with time constant parameter’ curve in Origin 2018. Taken the results of this experiment, the active SpCas9 variant quantities in solution were calculated. It was also taken into consideration that SpCas9 has a one-fold turnover rate.

### Determining cleavage rate of WT, B-HF1 and B-evoSpCas9 variants *in vitro*

At first, two different solutions were made: (1) target site containing plasmid solution and (2) an SpCas9-sgRNA master mix. After mixing them (see below) the ratio of the target site containing plasmid and active protein was 1:2. Both solutions were diluted with the same cleavage buffer (final concentration: 20 mM HEPES pH 7.5, 200 mM KCl, 2mM MgCl_2_, 1 mM TCEP, 2% glycerol) and were pre-incubated at 37 °C before reaction. To trigger cleavage reaction, the target site containing plasmid solution was added to the SpCas9-sgRNA mixture. To terminate cleavage reaction the inactivation solution (final concentration: 0.2% SDS, 50 mM EDTA) was added to the reaction mix at 80 °C at different time points. In case of the WT SpCas9 protein the sampling points were between 2 and 30 seconds, while in case of the increased-fidelity SpCas9 variants they fell between 5 seconds and 2 hours. To determine sampling points precisely a digital chronometer was attached to the pipette which can record time points in an application developed by us. This precise time determination was only necessary in the case of WT SpCas9 due to the fast reaction rate. Samples were then ran on a 0.8% agarose gel. Following densitometry (GelQuantNET, BiochemLabSolutions.com), the ratio of intact plasmid and total DNA was calculated for each sample. These values were plotted and fitted on a ‘One-phase exponential decay function with time constant parameter’ curve in Origin 2018. Experiments were performed in triplicates. All fitted curves are available in Supplementary Figure 6, the k values are available in Supplementary Table 8.

### Cell culturing and transfection

Cells employed in the studies are HEK293 (Gibco 293-H cells), GM08207 (Coriell Cell Repositories, Simian virus 40-transformed XP-D fibroblast), N2a.dd-EGFP (a neuro-2a mouse neuroblastoma cell line developed by us containing a single integrated copy of an EGFP-DHFR[DD] [EGFP-folA dihydrofolate reductase destabilization domain] fusion protein coding cassette originating from a donor plasmid with 1,000 bp long homology arms to the *Prnp* gene driven by the *Prnp* promoter (*Prnp.*HA-EGFP-DHFR[DD]), N2a.EGFP and HEK-293.EGFP (both cell lines containing a single integrated copy of an EGFP cassette driven by the *Prnp* promoter)^33^ cells. Cells were grown at 37 °C in a humidified atmosphere of 5% CO_2_ in high glucose Dulbecco’s Modified Eagle medium (DMEM) supplemented with 10% heat inactivated fetal bovine serum, 4 mM L-glutamine (Gibco), 100 units/ml penicillin and 100 μg/ml streptomycin. Cells were passaged up to 20 times (washed with PBS, detached from the plate with 0.05% Trypsin-EDTA and replated). After 20 passages, cells were discarded. Cell lines were not authenticated as they were obtained directly from a certified repository or cloned from those cell lines. Cells were tested for mycoplasma contamination.

Cells were plated in case of each cell line one day prior to transfection in 48-well plates at a density of approximately 2.5-3 x 10^4^ cells/well. Cells were co-transfected with two types of plasmids: SpCas9 variant expression plasmid (137 ng) and sgRNA and mCherry coding plasmid (97 ng) using 1 µl TurboFect reagent according to the manufacturer’s protocol. For negative control experiments either deadSpCas9 plasmid was co-transfected with a targeting sgRNA plasmid, or active SpCas9 variant with a non-targeting sgRNA plasmid. Transfection efficacy was calculated via mCherry expressing cells. Transfections were performed in triplicates. Transfected cells were analysed ∼96 h post-transfection by flow cytometry and genomic DNA was purified according to the Puregene DNA Purification protocol (Gentra systems).

### Plasmid and ribonucleoprotein electroporation

Briefly, 2 x 10^5^ cells were resuspended in transfection solution (see below) and mixed with 666 ng of SpCas9 variant expression plasmid and 334 ng of sgRNA and mCherry coding plasmid. In the case of GUIDE-seq experiments an additional 30 pmol dsODN (according to the original GUIDE-seq protocol^20^) was added to the mixture. For negative control experiments either a deadSpCas9 plasmid was co-transfected with a targeting sgRNA plasmid, or an active SpCas9 variant with a non-targeting sgRNA plasmid. Nucleofections were performed in the case of HEK293, GM08207 and HEK-293.EGFP cell lines using the CM-130 program on a Lonza 4-D Nucleofector instrument on strip, either with 20 µl SF solution according to the manufacturer’s protocol, or with 20 µl homemade nucleofection solution as described in Vriend et al.^76^. Transfection efficacy was calculated via mCherry expression. Unless noted otherwise, transfected cells were analysed ∼96 h post-transfection by flow cytometry followed by genomic DNA purification according to the Puregene DNA Purification protocol (Gentra systems) and downstream applications such as on-target amplicon PCR in three technical replicates.

In the case of EGFP 43, *FANCF* site 2 and *VEGFA* site 2 WT SpCas9 and SpCas9-HF1 GUIDE-seq experiments the electroporation was done as follows. Briefly, 2 x 10^6^ HEK293.EGFP or HEK293 cells were resuspended with 3 µg of SpCas9 variant expressing plasmid, 1.5 µg of mCherry and sgRNA coding plasmid and 100 pmol of the dsODN mixed together with 100 µl homemade nucleofection solution as described in Vriend et al.^76^. The mixture was electroporated using Nucleofector 2b (Lonza) with A23 program and 2 mm electroporation cuvettes.

*VEGFA* site 2 B-evo dRNA experiments were based on Rose et al.^64^. *VEGFA* sgRNA2 OT1 dRNA3 was used as follows: 1 x 10^6^ HEK293 cells were resuspended in 100 µl SF solution and mixed with 2.5 µg of B-evoSpCas9 expression plasmid and in case of dRNA 1:1 ratio: 1250 ng of dRNA3 and mCherry coding plasmid and 1250 ng of *VEGFA* site 2 sgRNA and mCherry coding plasmid, and in case of dRNA 6:1 ratio: 3000 ng of dRNA3 and mCherry coding plasmid and 500 ng of *VEGFA* site 2 sgRNA and mCherry coding plasmid. An additional 150 pmol GUIDE-seq dsODN was added to the mixture. Nucleofections were performed using the CM-130 program on a Lonza 4-D Nucleofector instrument in cuvettes according to the manufacturer’s protocol. Transfected cells were analysed ∼48 h post-transfection by flow cytometry. EGFP 43 WT, e- and SpCas9-HF1, *FANCF* site 2 WT, e-plus and HF1-plus and *VEGFA* site 2 WT and SpCas9-HF1 experiments are also described in Kulcsár et al.^21^.

In the case of RNP experiments with *VEGFA* site 3 HypaR-SpCas9 RNP, *VEGFA* site 2 B-evo SpCas9 RNP, 2 x 10^5^ HEK293 cells were transfected with 40 pmol SpCas9 and 48 pmol sgRNA (*VEGFA* site 3 HypaR-SpCas9 RNP 20 pmol SpCas9 and 24 pmol sgRNA), which was complexed in Cas9 storage buffer (20 mM HEPES pH 7.5, 200 mM KCl, 1 mM DTT and 10% glycerol) for 15 minutes at RT. 30 pmol of the dsODN was mixed with 20 µl SF solution to the RNP complex and electroporated using the CM-130 program on a Lonza 4-D Nucleofector instrument on strip. In case of *VEGFA* site 2 B-evo SpCas9 RNP, transfected cells were analysed ∼24 h post-transfection by flow cytometry. In the case of RNP experiments with EGFP 43 B-evo SpCas9 RNP, *FANCF* site 2 B-evo SpCas9 RNP, 2 x 10^6^ HEK293 or HEK293.EGFP cells were transfected with 100 pmol SpCas9 and 120 pmol sgRNA, which was complexed in Cas9 storage buffer (20 mM HEPES pH 7.5, 200 mM KCl, 1 mM DTT and 10% glycerol) for 15 minutes at RT. 100 pmol of the dsODN was mixed together with 100 µl homemade nucleofection solution to the RNP complex and electroporated using Nucleofector 2b (Lonza) with A23 program and 2 mm electroporation cuvettes.

### Flow cytometry

Flow cytometry analyses were carried out on an Attune NxT Acoustic Focusing Cytometer (Applied Biosystems). For data analysis Attune NxT Software v.2.7.0 was used. Viable single cells were gated based on side and forward light-scatter parameters and a total of 5,000 to 10,000 viable single cell events were acquired in all experiments. The GFP fluorescence signal was detected using the 488 nm diode laser for excitation and the 530/30 nm filter for emission, the mCherry fluorescent signal was detected using the 488 nm diode laser for excitation and a 640LP filter for emission or using the 561 nm diode laser for excitation and a 620/15 nm filter for emission. For detailed flow cytometry gating information see Supplementary Figure 1.

### EGFP disruption assay

EGFP disruption experiments were conducted in N2a.EGFP cells for the on-target screen (see details below), and in N2a.dd-EGFP cells for the mismatch screen with. Raw data of the EGFP disruption experiments are available in Supplementary Table 2, processed data of EGFP disruption experiments are available in Supplementary Table 3, heatmap data are available in Supplementary Table 4.

Background EGFP loss was determined for each experiment using co-transfection of dead SpCas9 expression plasmid and different targeting sgRNA and mCherry coding plasmids. EGFP disruption values were calculated as follows: the average EGFP background loss from dead SpCas9 control transfections made in the same experiment was subtracted from each individual treatment in that experiment and the mean values and the standard deviation (SD) were calculated from them. Results were normalized to the WT SpCas9 data from the same experiment.

**On-target activity** was measured in <u>N2a.EGFP</u> cell line. Cells were co-transfected with two types of plasmids: SpCas9 variant expression plasmid (137 ng) and sgRNA and mCherry coding plasmid (97 ng) using 1 µl TurboFect reagent per well in 48-well plates. Transfected cells were analysed ∼96 h post-transfection by flow cytometry. In this cell line the EGFP disruption level is not saturated, this way this assay is a more sensitive reporter of the intrinsic activities of these nucleases compared to N2a.dd-EGFP cell line.

In the case of **mismatch screens** <u>N2a.dd-EGFP</u> cells were co-transfected with two types of plasmids: with SpCas9 variant expression plasmid (137 ng) and a mix of 3 sgRNAs in which one nucleotide position was mismatched to the target using all 3 possible bases and mCherry coding plasmid (3 x∼33.3 ng = 97 ng) using 1 µl TurboFect reagent per well in 48-well plates. TMP (trimethoprim; 1 µM final concentration) was added to the media ∼48 h before FACS analysis. Transfected cells were analysed ∼96 h post-transfection by flow cytometry. Some of the data have also been shown in Kulcsár et al.^21^. The 4-day post-transfection results with this cell line show a close to saturated level, this way it is a good reporter system for seeing the full spectrum of off-target activities.

### Processing data from the study of Kim et al

Data from Kim et al.^34^ in Figure 4c-g were processed as follows. In case of the on-target screen, we selected those targets that were interrogated with perfectly matching tRNA-N_20_ protospacers (6,481 target sites) to avoid 5’ mismatched sgRNAs, then we excluded those targets that either lack data for any of the nucleases or were cleaved by the WT SpCas9 with lower than 15% indel occurrence.

In case of the mismatch screen, we processed the data as follows. We calculated the average of the on-target modification rates normalized to the corresponding WT values from the parallel experiments, and for further processing, we selected data from only those off-targets and IFNs, where the corresponding average on-target values normalized to the WT were at least 0.20 measured on day 4. We considered only the one base mismatching targets: off-targets with every possible one base mismatch for all positions, i.e., 60 data points per sgRNA, and for all the 30 sgRNAs per SpCas9 variant (i.e., 1,800 datapoints overall). The average of the modification (indel) percentages of the 60 off-target values for each sgRNA and IFN pair were calculated and normalized to the corresponding on-target value of the SpCas9 variants on day 7. These are presented along with the day 7 on-target data in the heatmap in Figure 4f. For detailed information see Supplementary Table 7.

### On-target heatmaps

The algorithms for ordering rows and columns on the on-target heatmaps is the following: After subtracting the background, normalized on-target values were calculated by dividing them with the WT value and then rounding them to two decimals. Values that were below zero were rounded to zero. Values lower than 0.20 were regarded as no cleavage. Heatmaps were ordered as follows: (i) IFNs were ordered according to how many targets they could cleave. When the number of cleaved targets was the same for multiple IFNs, they were ordered according to their average normalized on-target activity. (ii) Targets were ordered based on the number of IFNs that can cleave them, taking it into consideration to minimize the number of outliers. On each heatmap, a bold line shows the threshold between cleaved and non-cleaved datapoints, and outliers are clearly indicated.

### Binary classification

G-mean is the squared root of the product of the sensitivity and specificity that was calculated for the entire on-target heatmap for the cleaved and non-cleaved groups, where the bold line indicates where the cleavability law predicts the border between cleaved (≥0.20) and non-cleaved (<0.20) values. For G-mean calculation data (confusion matrix, sensitivity and specificity) see Supplementary Table 4. ROC curves are graphs that plot a model’s false-positive rate against its true-positive rate across a range of classification thresholds. ROC curves were generated for individual columns of the on-target heatmaps representing the normalized on-target activity values for a variant to assess how accurately the cleavage rule ordered its targets into cleaved and non-cleaved classes. For ROC curve and AUC calculation data see Supplementary Table 4.

### ssODN repair of *ERCC2* exon22 R683W (2047C>T) mutation

Donor ssODN for GM08207 cell line *ERCC2* exon22 R683W (2047C>T) mutation repair was designed to have the wild type base and a silent mutation (to identify the repair outcome). The 90Lnt long ssODN was centered at the desired mutations (Figure 6a and Supplementary Table 1: PCR primers/ERCC2 90nt + marked primer). Briefly, 2 x 10^5^ GM08207 cells were resuspended in 20 µl homemade nucleofection solution as described in Vriend et al.^76^ and mixed with 666 ng of SpCas9 variant expression plasmid and 334 ng of sgRNA and mCherry coding plasmid and 2 µl of 100 µM ssODN donor. Nucleofections were performed using the CM-130 program on a Lonza 4-D Nucleofector instrument. Cells were plated in 48-well plates containing 0.5 ml of completed DMEM and 2 µM M3814 HDR enhancer^62^ (which was a kind gift from Stephan Riesenberg) per well. After two days media was changed to fresh completed DMEM. Transfections were performed in triplicates. For negative control experiments deadSpCas9 plasmid was co-transfected with the targeting sgRNA plasmid. Transfected cells were analysed ∼96 h post-transfection by flow cytometry and genomic DNA was purified according to the Puregene DNA Purification protocol (Gentra systems). For NGS data information see Supplementary Table 5.

### Indel analysis by next-generation sequencing (NGS)

Amplicons for deep sequencing were generated using two rounds of PCR to attach Illumina handles. The 1^st^ step PCR primers used to amplify target genomic sequences are listed in Supplementary Table 1: PCR primers. PCR was done in a S1000 Thermal Cycler (Bio-Rad) or PCRmax Alpha AC2 Thermal Cycler using the by Q5 high-fidelity polymerase with supplied Q5 buffer (in case of VEGFA site 2 amplicon together with Q5 High GC enhancer) and 150 ng of genomic DNA in a total volume of 25 μl. The thermal cycling profile of the PCR was: 98 °C 30 sec; 35 x (denaturation: 98 °C 20 sec; annealing: see Supplementary Table 1: PCR primer, 30 sec; elongation: 72 °C, see Supplementary Table 1: PCR primer); 72°C 5 min. i5 and i7 Illumina adapters were added in a second PCR reaction using Q5 high-fidelity polymerase with supplied Q5 buffer (in case of VEGFA site 2 amplicon together with Q5 High GC enhancer) and 1 µl of first step PCR product in total volume of 25 μl. The thermal cycling profile of the PCR was: 98 °C 30 sec; 35 x (98 °C 20 sec, 67 °C 30 sec, 72 °C 20 sec); 72°C 5 min. Amplicons were purified by agarose gel electrophoresis. Samples were quantified with Qubit dsDNA HS Assay kit and pooled. Double-indexed libraries were sequenced on a MiSeq, MiniSeq or NextSeq (Illumina) giving paired-end sequences of 2 x 150 bp or 2 x 250 bp, it was performed by ATGandCo or Deltabio Ltd. Reads were aligned to the reference sequence using BBMap. Indels were counted computationally amongst the aligned reads that matched at least 75% of the first 20 bp of the reference amplicon. Indels without mismatches were searched starting at ±2 bp around the cut site. For each sample, the indel frequency was determined as (number of reads with an indel) / (number of total reads). The 15 bp long centre fragment of the GUIDE-seq dsODN sequence (“gttgtcatatgttaa” / “ttaacatatgacaac”) was counted in the aligned reads to measure dsODN on-target tag integration for GUIDE-seq experiments. The ssDNA repair was determined as (number of reads with desired edit) / (number of total reads). Results can be found in Supplementary Table 2. The following software were used: BBMap 38.08, samtools 1.8, BioPython 1.71, PySam 0.13. For NGS data information see Supplementary Table 5 and NGS sequencing data are deposited at NCBI Sequence Read Archive and will be available upon publication.

### GUIDE-seq

GUIDE-seq relies on the integration of a short dsODN tag into DNA breaks, therefore after the genomic DNA purification, dsODN tag integration and efficient indel formation was verified in the on-target site by NGS. In the next step genomic DNA was sheared with BioraptorPlus (Diagenode) to 550 bp in average. Sample libraries were assembled as previously described^20^ and sequenced on Illumina MiSeq or MiniSeq instrument by ATGandCo or Deltabio Ltd. Data were analysed using open-source guideseq software (version 1.1)^77^. Consolidated reads were mapped to the human reference genome GrCh37 supplemented with the integrated EGFP sequence. Upon identification of the genomic regions integrating double-stranded oligodeoxynucleotide (dsODNs) in aligned data, off-target sites were retained if at most seven mismatches against the target were present and if absent in the background controls. Visualization of aligned off-target sites are provided as a color-coded sequence grid. Summarized results can be found in Supplementary Table 6 and GUIDE-seq sequencing data are deposited at NCBI Sequence Read Archive and will be available upon publication.

### Western blot

N2a.dd-EGFP cells were cultured on 48-well plates and were transfected as described above in the EGFP disruption assay section. Four days post-transfection, 9 parallel samples corresponding to each SpCas9 variant transfected were washed with PBS, then trypsinized and mixed, and were analysed for transfection efficiency via mCherry fluorescence level by using flow cytometry. The cells from the mixtures were centrifuged at 200 rcf for 5 min at 4 °C. Pellets were resuspended in ice cold Harlow buffer (50 mM Hepes pH 7.5; 0.2 mM EDTA; 10 mM NaF; 0.5% NP40; 250 mM NaCl; Protease Inhibitor Cocktail 1:100; Calpain inhibitor 1:100; 1 mM DTT) and lysed for 20-30 min on ice. The cell lysates were centrifuged at 19,000 rcf for 10 min. The supernatants were transferred into new tubes and total protein concentrations were measured by the Bradford protein assay. Before SDS gel loading, samples were boiled in Protein Loading Dye for 10 min at 95 °C. Proteins were separated by SDS-PAGE using 7.5% polyacrylamide gels and were transferred to a PVDF membrane, using a wet blotting system (Bio-Rad). Membranes were blocked by 5% non-fat milk in Tris buffered saline with Tween20 (TBST) (blocking buffer) for 2 h. Blots were incubated with primary antibodies [anti-FLAG (F1804, Sigma) at 1:1,000 dilution; anti-β-actin (A1978, Sigma) at 1:4,000 dilution in blocking buffer] overnight at 4 °C. The next day, after washing steps in TBST, the membranes were incubated for 1 h with HRP-conjugated secondary anti-mouse antibody 1:20,000 (715-035-151, Jackson ImmunoResearch) in blocking buffer. The signal from detected proteins was visualized by ECL (Pierce ECL Western Blotting Substrate, Thermo Scientific) using a CCD camera (Bio-Rad ChemiDoc MP).

### Statistics

Differences between SpCas9 variants were tested by using either two-tailed paired-samples Student’s t-test (Fig. 6b SpCas9-NG/SpCas9-NG-L847R-V1015R) or by using two-tailed Wilcoxon Signed Ranks test (Fig. 6b HypaR/HypaR-L847R-V1015R, xCas9/xCas9-L847R- V1015R) in the cases where differences did not meet the assumptions of Paired t-test. Differences between groups were tested by using either two-tailed unpaired Student’s t-test with Welch’s correction (Fig. 5c B-SpCas9-HF1) or by using two-tailed Mann-Whitney test (Fig. 5c B-evoSpCas9) in the cases where differences did not meet the assumptions of unpaired t-test. Differences between SpCas9 variants were tested by using RM one-way ANOVA and Dunnett’s multiple comparisons test with a single pooled variance (Fig. 4b) or by using RM one-way ANOVA, with the Geisser-Greenhouse correction and Dunnett’s multiple comparisons test with individual variances computed for each comparison (Fig. 6a) or (ii) Tukey’s multiple comparisons test with individual variances computed for each comparison (where the mean of each column was compared with the mean of every other columns: Fig. 2b, 4d, Supplementary Fig. 2i) in the cases where sphericity did not meet the assumptions of RM one-way ANOVA. Differences between more than two groups were tested by using Kruskal-Wallis test (Fig. 2e, 4g, Supplementary Fig. 3d). Normality of data and of differences was tested by Shapiro-Wilk normality test. Statistical tests were performed using GraphPad Prism 9 on data including all parallel sample points. Test results are shown in Supplementary Table 9.

## Supplementary Figures and Tables

**Supplementary Figure 1.**
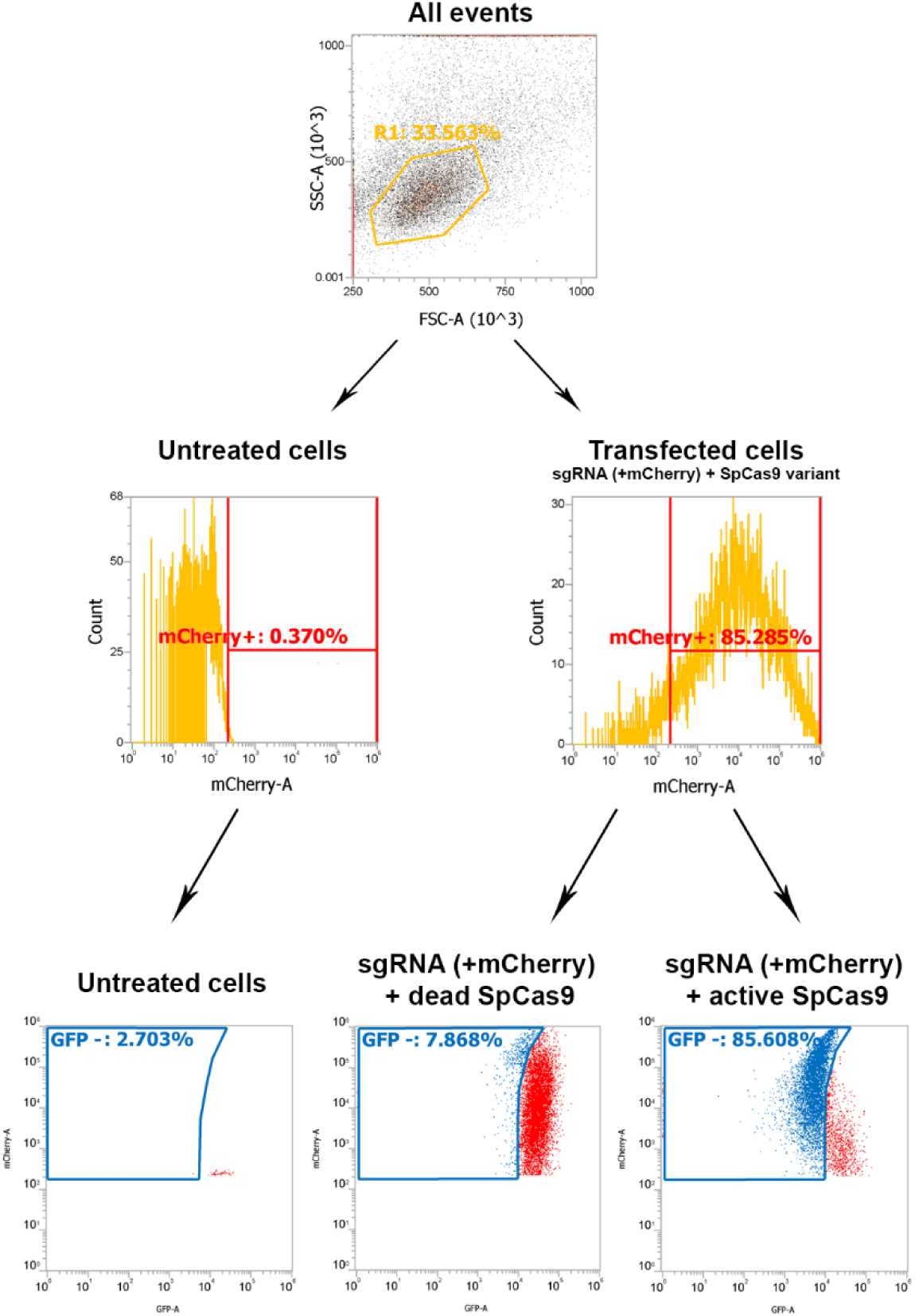
Flow cytometry gating example is shown for experiments with N2a.dd-EGFP, N2a.EGFP, HEK-293.EGFP, HEK293 and GM08207 cell lines. Live, single cells were gated by FSC and SSC parameters (upper panel - R1). Middle panels show mCherry FACS histograms (for R1 gated cells). Middle left panel shows an example for untreated (non-transfected) cells, middle right panel for transfected cells. In the case of HEK293 and GM08207 cells only the percentage of mCherry positive cells was determined by flow cytometry. In the case of N2a.dd-EGFP, N2a.EGFP and HEK-293.EGFP cells GFP fluorescence was measured in the mCherry positive population (lower panels). The lower panels show the gated GFP negative cells (GFP loss in EGFP disruption) in the case of untreated cells (lower left panel), cells transfected with sgRNA and mCherry coding plasmid and dead SpCas9 coding plasmid (lower middle panel) and cells that were transfected with sgRNA and mCherry coding plasmid and active SpCas9 variant coding plasmid (lower right panel).

**Supplementary Figure 2.**
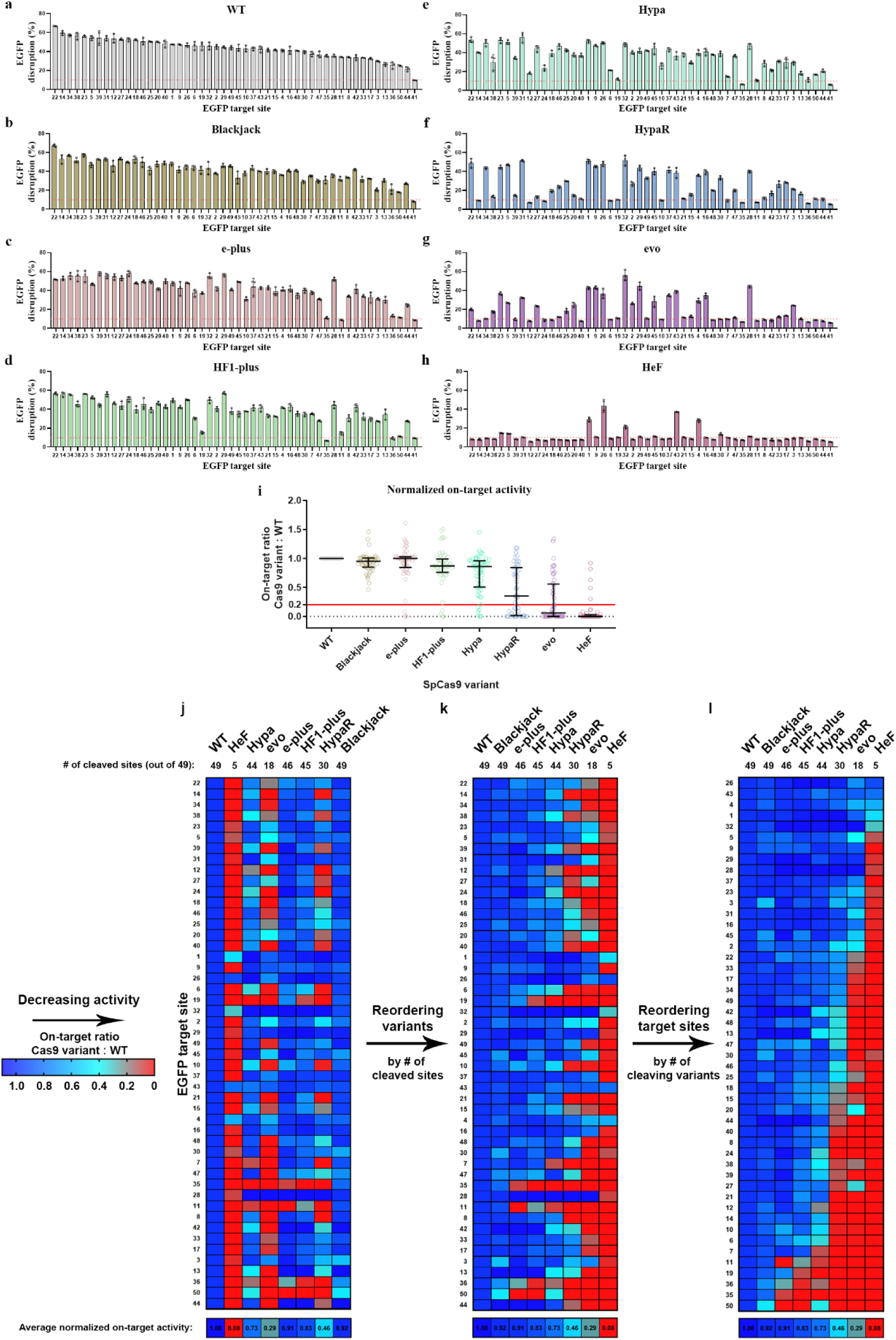
The cleavage rule becomes apparent by rearranging the normalized disruption data. **a-h,** On-target EGFP disruption data of (**a**) WT SpCas9 and (**b-h**) seven IFN variants, as indicated in the panels, on 50 EGFP target sites. Means and SD are shown; n=3 biologically independent samples (overlaid as white circles); level of background EGFP loss is indicated by the red dashed line (average of the percentage of dead SpCas9 controls from all target sites). The targets are arranged in a descending order based on WT disruption values in all panels. EGFP site 41 was not even cleaved by the WT SpCas9, and therefore it is not included in any downstream figures. This accounts for the difference in the number of targets (50 vs. 49) between panels (**a-h**) and other figures. **i,** On-target disruption activities normalized to the WT disruption values of different SpCas9 variants presented on a scatter dot plot. The sample points correspond to data presented in panels (**a-h**). Continuous red line indicates 0.20 normalized disruption activity, under which we consider the IFNs not to be active on a given target. The median and interquartile range are shown; data points are plotted as open circles representing the mean of biologically independent triplicates. Statistical significance was assessed by using RM one-way ANOVA and shown in Supplementary Table 9. **j-l,** Heatmaps show the normalized EGFP disruption activity of SpCas9 nucleases with perfectly matching sgRNAs. **j**, The targets are arranged in a descending order based on WT disruption values, like in panels (**a-h**). **k,** IFNs were reordered according to how many targets they could cleave, and **l**, targets were reordered based on how many IFNs could cleave them (same pattern as in Supplementary Figure 3a). Reordering highlights the rankings amongst targets and IFNs and reveal the cleavage rule; if a target can be cleaved by a nuclease variant, then it will be cleaved by all variants with a higher activity/lower fidelity rank. **a**-**l**, Target sequences, raw, processed and heatmap disruption data and statistical details are reported in Supplementary Table 1-4 and 9.

**Supplementary Figure 3.**
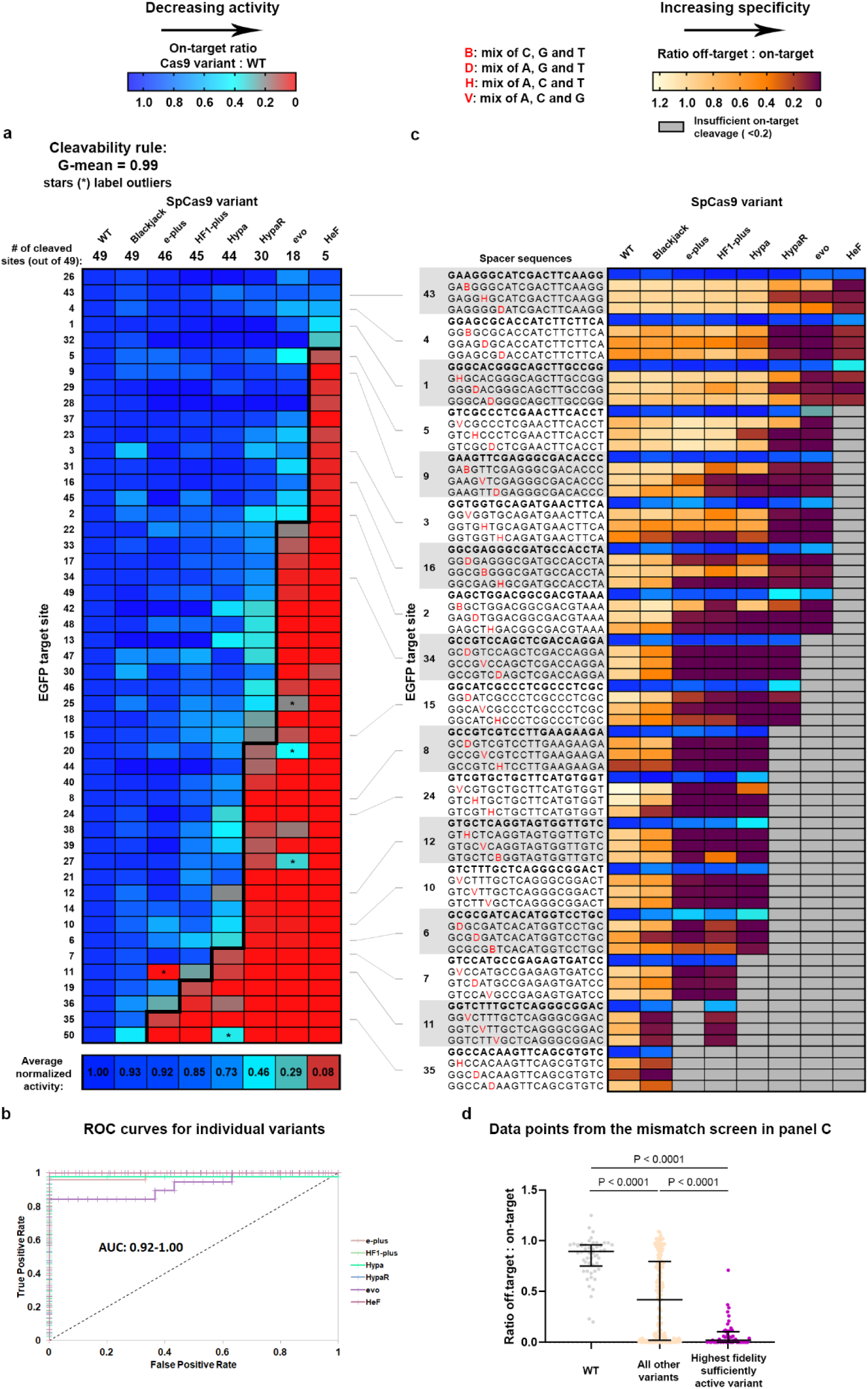
Cleavage rule enables the selection of the most optimal increased fidelity nuclease. Heatmaps show normalized EGFP disruption activities of SpCas9 nucleases with (**a**) perfectly matching and (**c**) partially mismatching 20G-sgRNAs. **a,** The bold line indicates the dividing line defined by the cleavage rule between classes of cleaved and not-cleaved values. The G-mean value indicates how well the data points above and below the bold line correspond to cleaved and not-cleaved (<0.20 activity normalized to WT) experimental values. Outliers have been confirmed in repeated experiments. **b,** The ROC curves demonstrate that the order of the target sequences, determined by the cleavage rule, competently separates the classes of cleaved and not-cleaved normalized disruption values of each individual variant from panel (**a**). This indicates that IFNs universally perceive the target contributions that primarily determine whether they cleave a particular target or not. **c**, Targets from higher ranks (cleavable by many IFNs) require higher fidelity nucleases, while targets from lower ranks (cleavable by few IFNs) require lower fidelity nucleases for editing them with both high efficiency and high specificity. **d,** Matching IFNs and targets further increases the specificity of editing. The median and interquartile range of data points that are selected from panel (**c**) is presented as indicated; n=54, 189, 54, respectively. Dots are shown for each variant with each mismatching spacer position, where the normalized on-target activity exceeded 70%. **a-d,** Target sequences, raw and processed disruption data and statistical details are reported in Supplementary Table 1-4 and 9.

**Supplementary Figure 4.**
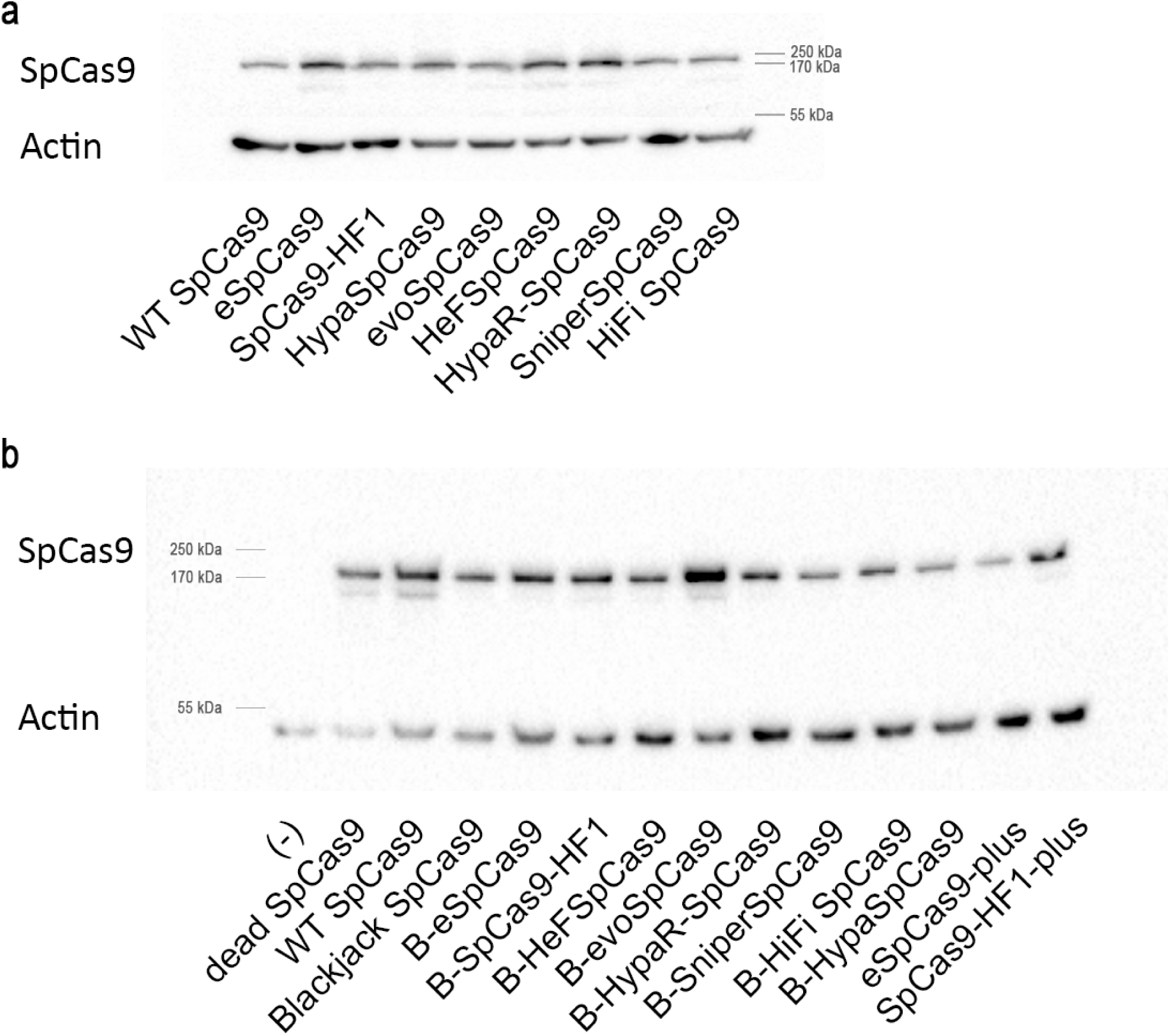
Immunoblot analysis of the expression levels of SpCas9 nuclease variants. SpCas9 variants (∼160 kDa) in cell lysates of reporter N2a.dd-EGFP cells transfected with the indicated nuclease constructs. β-actin (∼42 kDa) was used as loading control for total protein amounts analysed.

**Supplementary Figure 5.**
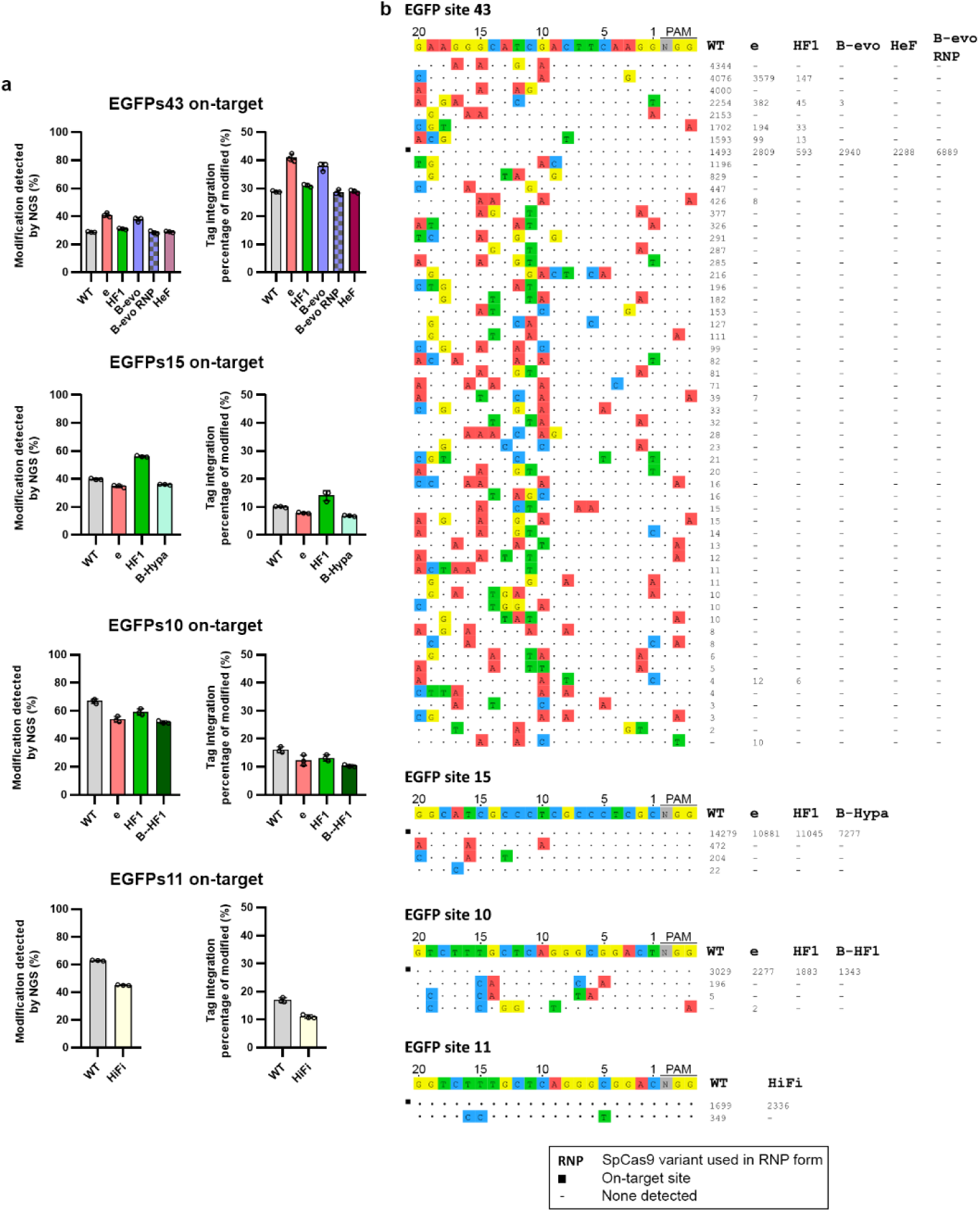
Details of the GUIDE-seq experiments presented in. Figure 3. Modifications at on-target and off-target cleavage sites of SpCas9 variants with sgRNAs targeting EGFP sites identified in GUIDE-seq experiments. **a**, The percentage of on-target genome modification (indel + tag integration) and the tag integration frequency of the modified cells analysed by NGS are presented in the bar charts. Means and SD are shown; nL=3 (overlaid as white circles). **b,** Read counts that give an approximate measure of cleavage frequency at a given sequence are shown; mismatched positions within the spacer or PAM are highlighted in different colours. (-) indicates zero reads, which means that off-target cleavage was not detected; black squares indicate the on-target sites. **a-b**, Data related to Figure 3 and 9. Target sequences, NGS and GUIDE-seq data are reported in Supplementary Tables 1, 5 and 6.

**Supplementary Figure 6.**
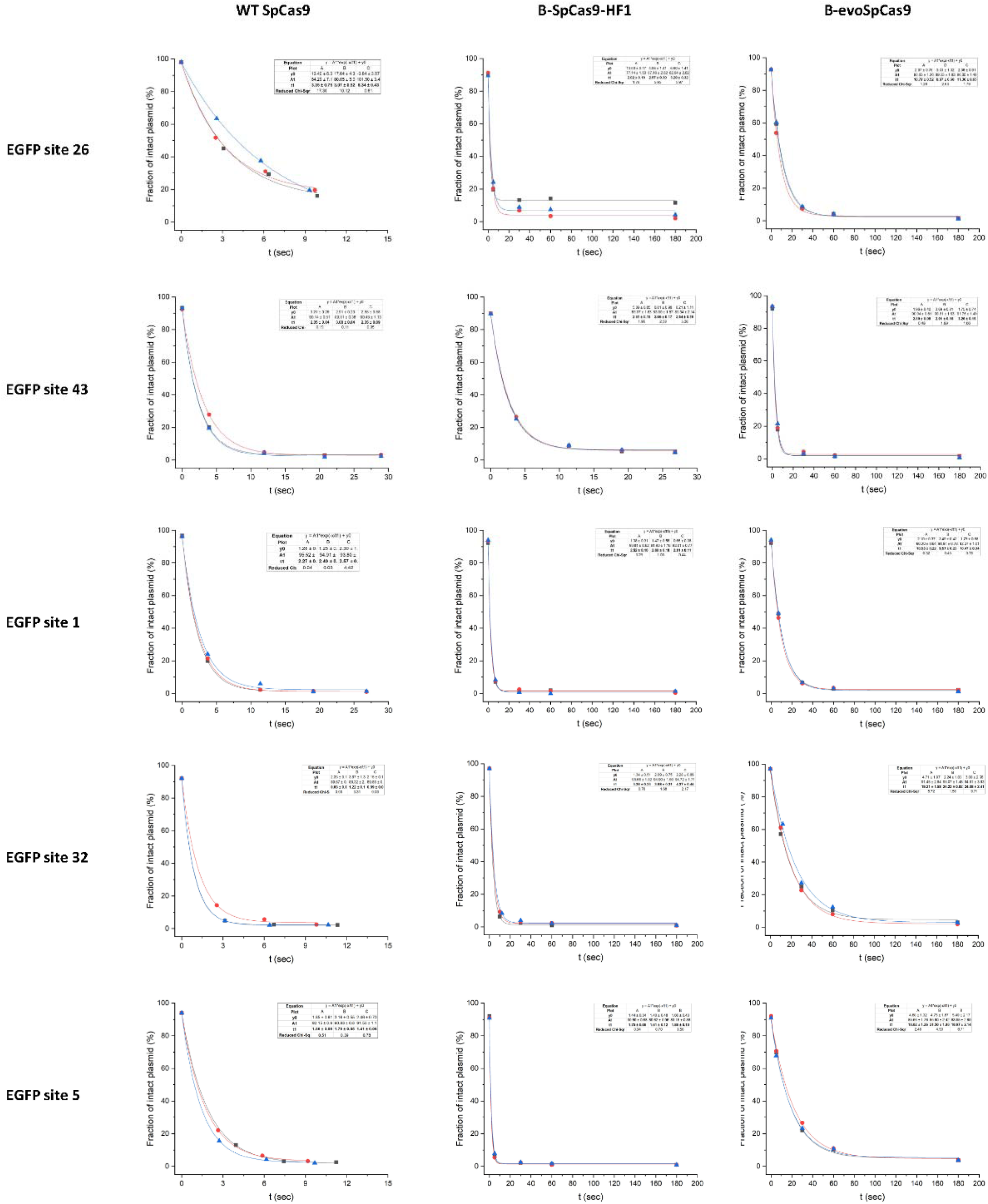
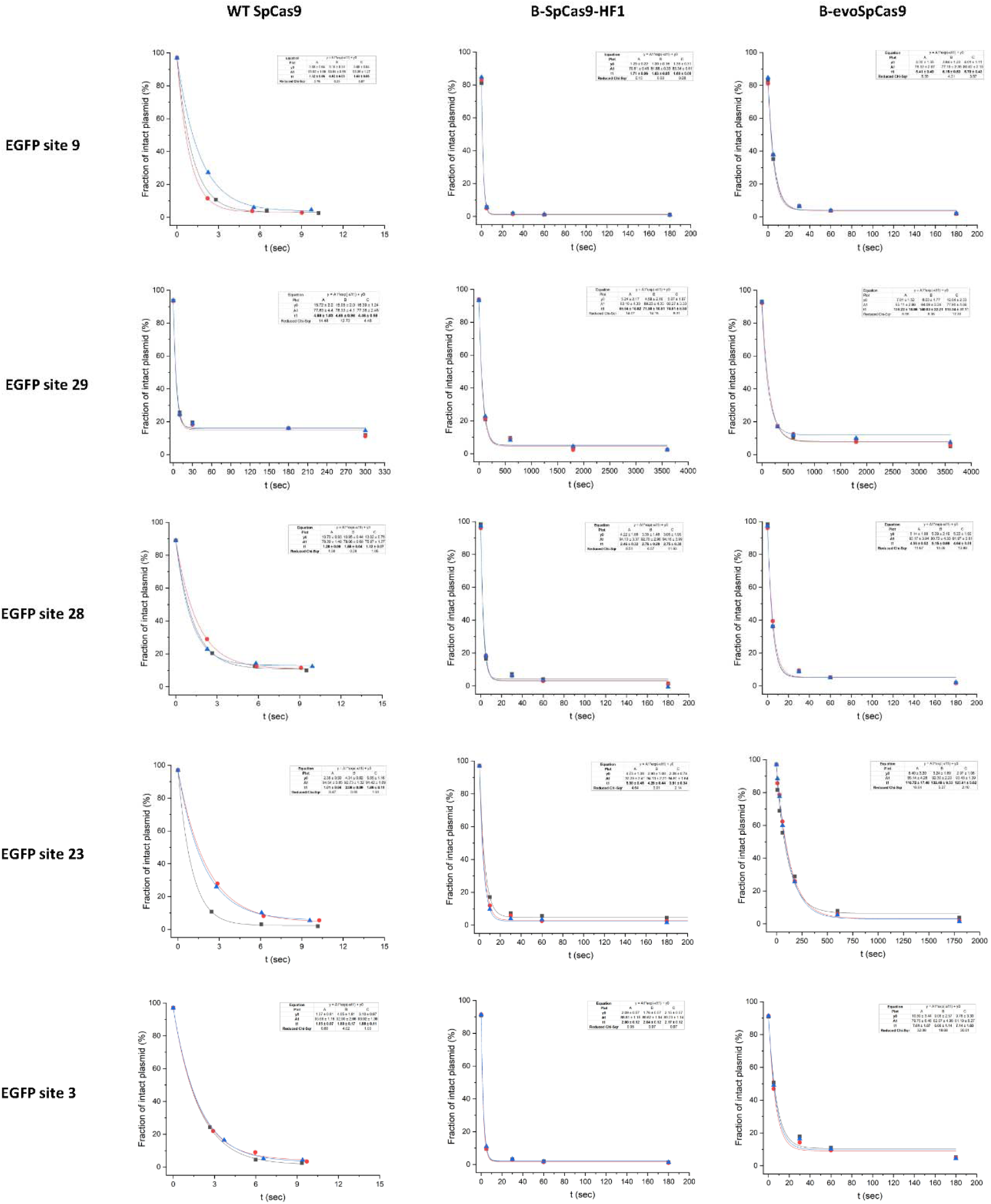
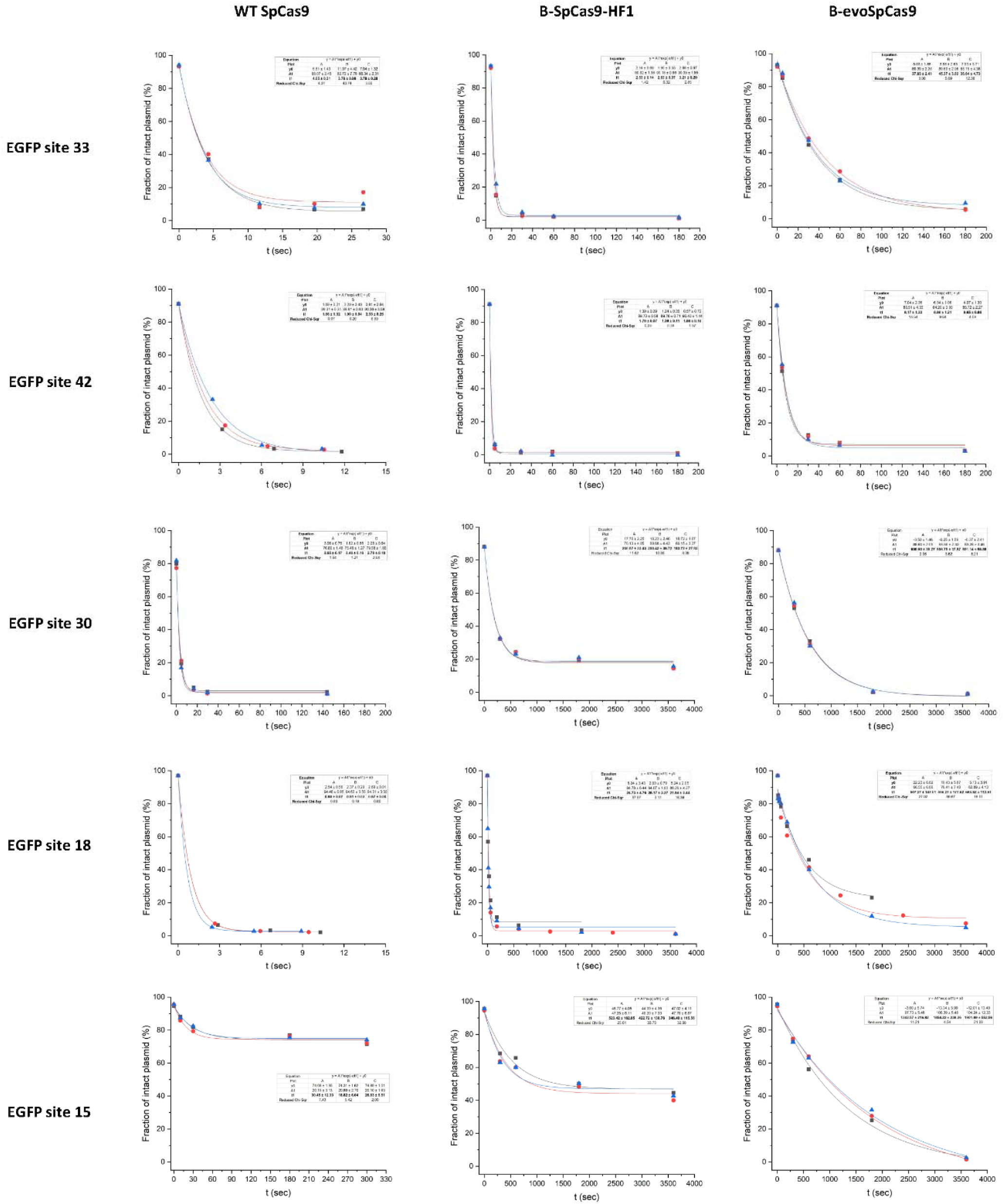
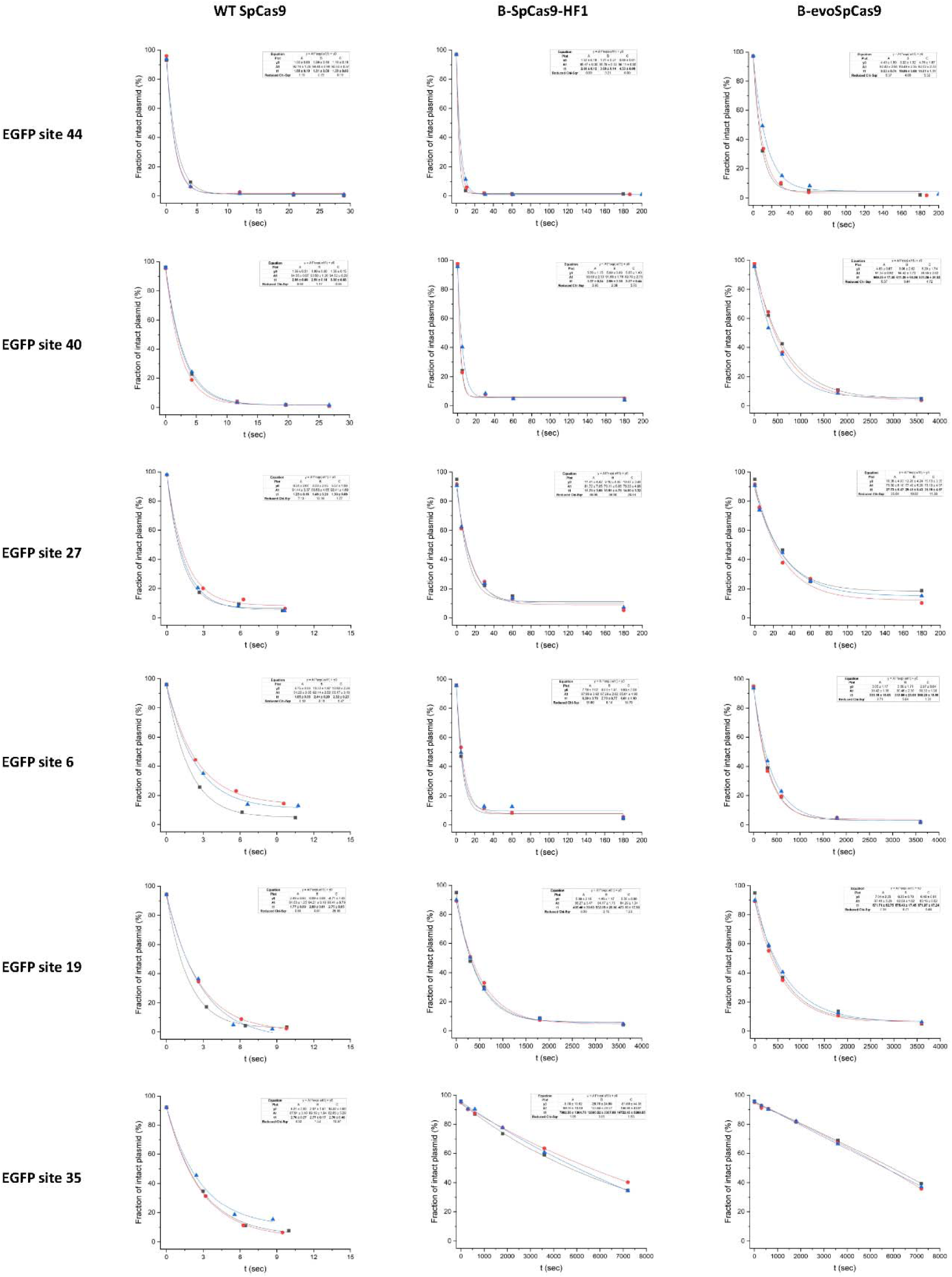
*In vitro* fitted curves on 21 EGFP target sites cleaved with WT, B- HF1 or B-evoSpCas9 variants. The experiments were conducted as follows: The RNP complex was added to circular plasmid DNA (bearing the EGFP target site complementary to the spacer of the sgRNA) and the fraction of intact plasmid was measured at different time points. 0 sec was determined by running a plasmid only control. Plots show values derived from the band intensities measured on agarose gel. Exponential curves were fitted to the fraction of intact plasmid measured during the time frame of the experiments, in the case of each replicate separately. The values and average k values were derived from these fitted curves. Data related to Figure 5. Summary of target and primer sequences and *in vitro* data are reported in Supplementary Table 1 and 8.

**Supplementary Figure 7.**
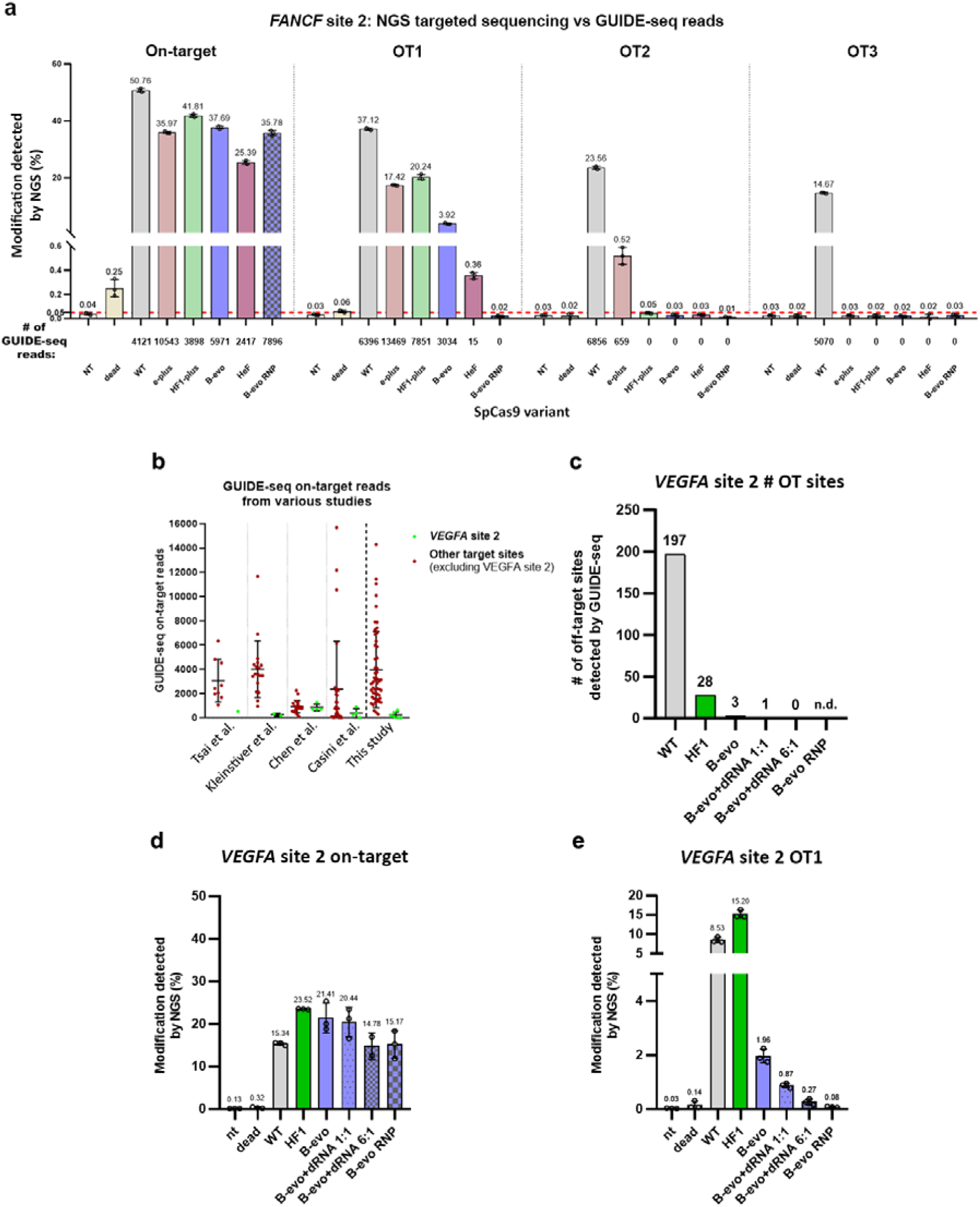
NGS confirms the editing of *VEGFA* site 2 without any off-target. **a,** GUIDE-seq and amplicon sequencing show comparable sensitivity for assessing off-target editing determined from the same *FANCF* site 2 samples for both methods. The percentage of genome modification (indel + tag integration) of on-target and of three off-target sites analysed by NGS are shown in the bar charts. Means and SD are shown; n=3 (overlaid as white circles), the red dashed line indicates the commonly estimated limit of detection. GUIDE-seq read counts that give an approximate measure of cleavage frequency at a given sequence are shown under the graph. **b,** Scatter plot shows that the mean of the on-target GUIDE-seq reads for all targets used in a given study (for details see Supplementary Table 6) is one of the highest in this study, stressing the significance of the editing without any off-targets demonstrated here. The on-target GUIDE-seq reads for *VEGFA* site 2 target are unusually low across all studies compared to most other target sites preventing a straightforward interpretation of the off-target reads. In the scatter dot plot means and SD are shown; data points are plotted as circles representing the on-target GUIDE-seq reads in the case of every tested target site in a given paper. **c-e,** NGS confirms the editing of *VEGFA* site 2 without any off-target. When there are only a few or no off-target GUIDE-seq reads, the results are not meaningful unless the number of on-target GUIDE-seq reads and the percentage of genome modification from the same sample, determined by NGS, are known. **c,** Bar chart showing the total number of off-target sites detected by GUIDE-seq. Due to the few on-target reads for *VEGFA* site 2, we performed amplicon sequencing using the samples of panel (**c**) to sequence the (**d**) on-target site and the (**e**) off-target site 1 (OT1), its most persistent off-target sequence. **d, e,** The percentage of genome modification (indel + tag integration) is shown in bar charts. Means and SD are shown; n=3 (overlaid as white circles). **c-e,** While on-target modification rates are comparable with the WT SpCas9 in all conditions in panel (**d**), OT1 reads in panel (**e**) are decreasing in accordance with the total number of off-target sites detected by GUIDE-seq (**c**) and with the GUIDE-seq reads in Supplementary Figure 9. The combination of B-evoSpCas9 with dRNA^64^ (B-evo+dRNA 6:1) diminishes off-target editing detected by GUIDE-seq although showing detectable OT1 modification by NGS as shown in panel (**c**) and Supplementary Figure 8c, and low on-target GUIDE-seq read counts (Supplementary Fig. 9). In panel (**c**), in case of B-evoSpCas9 in RNP form no GUIDE-seq read was detected (n.d.) even though it showed WT-like on-target cleavage (**d**) and tag integration (Supplementary Fig. 8b). B-evoSpCas9 in RNP form showed a modification rate on OT1 site, that is smaller than that of dead SpCas9, falling under the detection limit of NGS (**e**) and showed no tag integration (Supplementary Fig. 8c). **a-e,** Data related to Figures 3, 7-10 and Supplementary Figures 5, 8 and 9. Target sequences, NGS and GUIDE-seq data are reported in Supplementary Tables 1, 5 and 6.

**Supplementary Figure 8.**
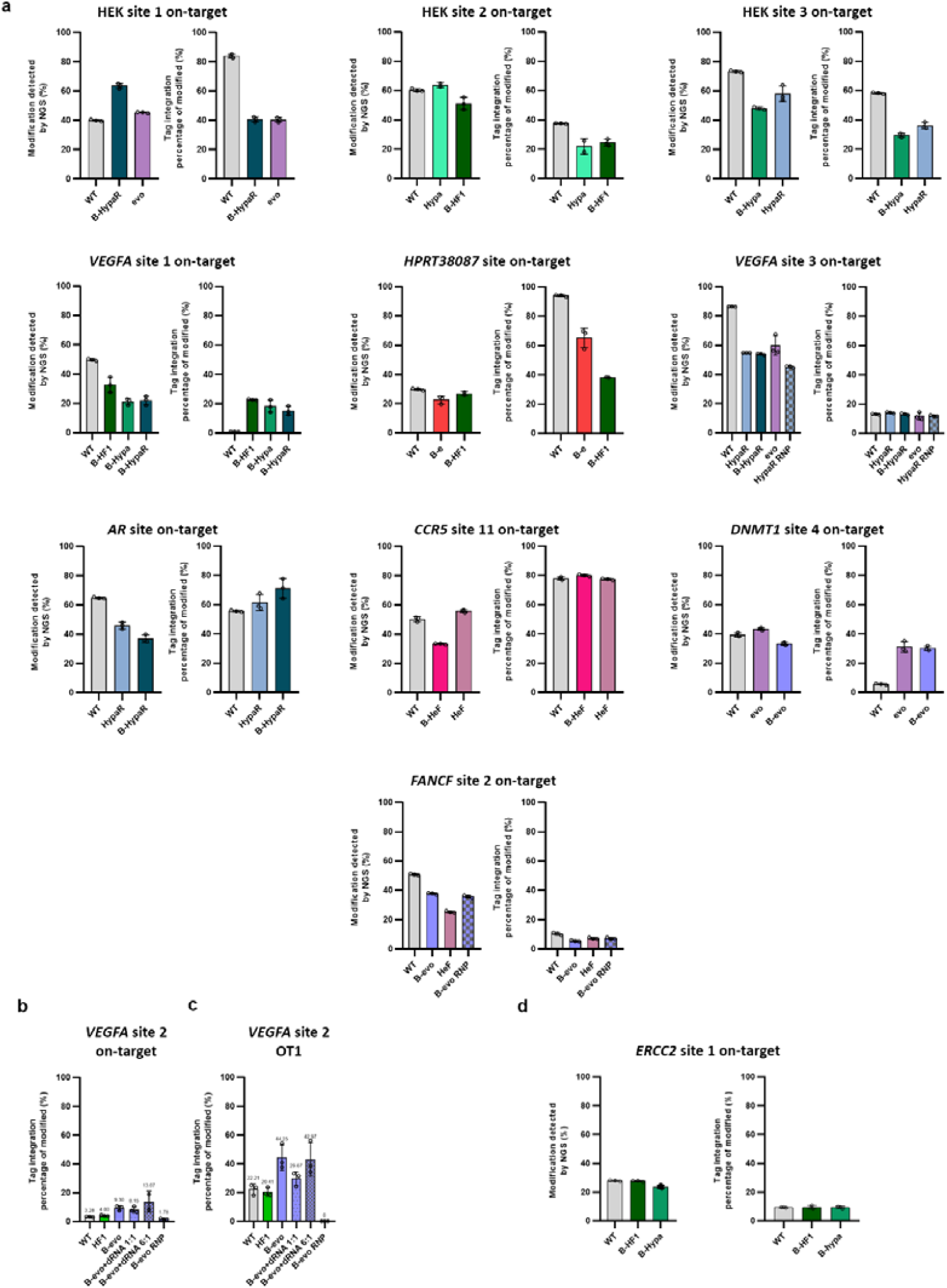
On-target modification rates and GUIDE-seq dsODN tag integration rates detected by NGS. **a-c,** The percentage of on-target genome modification (indel + tag integration) and the tag integration frequency of the modified cells are shown in bar charts, as indicated in the figure. Means and SD are shown; n=3 (overlaid as white circles). Data related to Figures 7, 8, 10 and Supplementary Figures 7, 9. Summary of NGS data is reported in Supplementary Table 5.

**Supplementary Figure 9.**
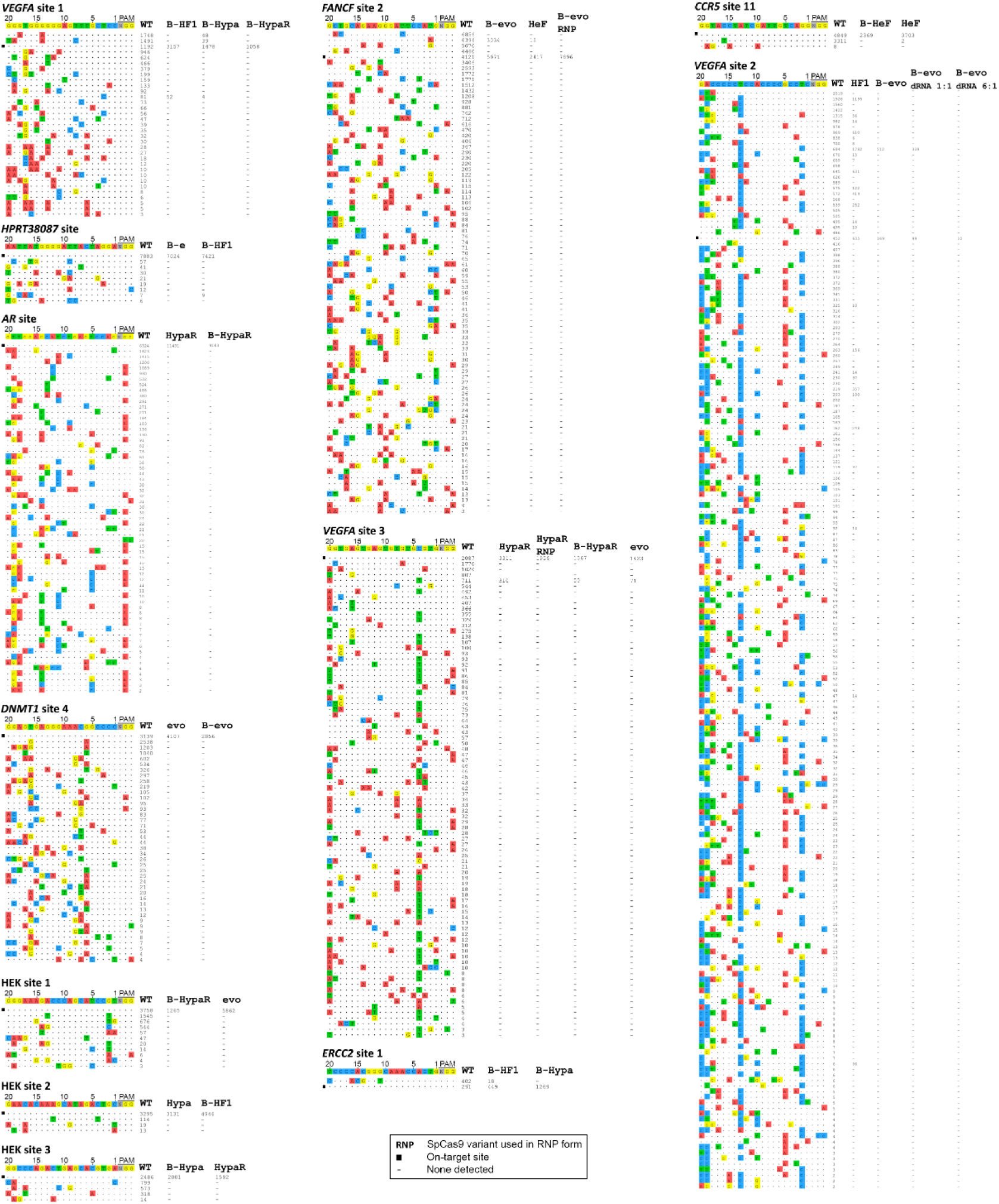
Genome-wide off-target detection of increased fidelity SpCas9 variants using GUIDE-seq. Off-target cleavage sites of SpCas9 variants identified by GUIDE-seq. Read counts that give an approximate measure of cleavage frequency at a given sequence are shown, mismatched positions within the spacer or the PAM are highlighted in different colours. (-) indicates zero reads, which means that off-target cleavage was not detected, black squares indicate on-target sites, data with no GUIDE-seq read are not shown (*VEGFA* site 2 B-evoSpCas9 in RNP form). Data related to Figures 7, 8, 10 and Supplementary Figures 7, 8. Summary of GUIDE-seq data is reported in Supplementary Table 6.

**Supplementary Table 10.** Summary of potential off-target sites identified by Cas-OFFinder^78^ in the reference human genome for the sgRNAs examined in this study by GUIDE-seq. Data related to Figure 9.

| site | mismatches to on-target site |  |  |  |  |  | total |
| --- | --- | --- | --- | --- | --- | --- | --- |
|  | 1 | 2 | 3 | 4 | 5 | 6 |  |
| EGFP site 43 | 0 | 0 | 5 | 88 | 827 | 5895 | 6815 |
| CCR5 site 11 | 0 | 1 | 5 | 125 | 1391 | 10111 | 11633 |
| VEGFA site 2 | 0 | 1 | 12 | 143 | 1519 | 10877 | 12552 |
| FANCF site 2 | 0 | 2 | 35 | 456 | 3905 | 17576 | 21974 |
| DNMT1 site 4 | 1 | 1 | 29 | 235 | 2000 | 13047 | 15313 |
| VEGFA site 3 | 0 | 2 | 17 | 251 | 1732 | 11335 | 13337 |
| HEK site 1 | 0 | 0 | 9 | 96 | 1065 | 7675 | 8845 |
| VEGFA site 1 | 0 | 1 | 11 | 88 | 857 | 6401 | 7358 |
| AR site | 1 | 4 | 35 | 281 | 2202 | 14064 | 16587 |
| HEK site 3 | 1 | 1 | 0 | 20 | 273 | 2807 | 3102 |
| ERCC2 site 1 | 1 | 17 | 383 | 6089 | 13536 | 35901 | 55927 |
| EGFP site 15 | 0 | 3 | 10 | 158 | 1377 | 9833 | 11381 |
| EGFP site 10 | 0 | 0 | 2 | 78 | 875 | 6790 | 7745 |
| HEK site 2 | 0 | 3 | 16 | 340 | 3153 | 13510 | 17022 |
| HPRT38087 site | 0 | 0 | 4 | 62 | 731 | 5649 | 6446 |
| EGFP site 11 | 0 | 0 | 6 | 69 | 1021 | 8733 | 9829 |

## Acknowledgements

We thank Ildikó Szűcsné Pulinka, Judit Szűcs, Vivien Karl, Lilla Burkus, Barbara Karsai, Judit Kálmán for their excellent laboratory assistance, Dorottya Simon, Antal Nyeste, Zsuzsa Bartos, Edit Szabó, György Várady for their valuable help. We thank Stephan Riesenberg for his valuable advice and providing the HDR enhancer^62^. We thank Dóra Bokor for proofreading the manuscript. This research was supported by grants K128188 and K134968 to E.W and PD134858 to P.I.K. from the Hungarian Scientific Research Fund (OTKA) and P.I.K. by 2018- 1.1.1-MKI-2018-00167 and by ÚNKP-20-5-SE-20 from the National Research, Development and Innovation Office. P. I. K. is a recipient of the János Bolyai Research Scholarship of the Hungarian Academy of Sciences (BO/764/20). S.L.K was supported by grant EFOP-3.6.3- VEKOP-16-2017-00009 from the Higher Education Institutional Excellence Program of the Semmelweis University.

## Author contributions

P.I.K. and E.W. conceived and designed experiments, interpreted the results, P.I.K., A.T., Z. L., E. T., V. V., R. Z., S. L. K., K. H. performed all experiments. P.I.K., A.T., E. T., Z. L., R. Z., S.

L. K analysed the data. P.I.K. and E.W. wrote the manuscript with input from all the authors.

